# What can we learn when fitting a simple telegraph model to a complex gene expression model?

**DOI:** 10.1101/2023.03.09.532005

**Authors:** Feng Jiao, Jing Li, Ting Liu, Yifeng Zhu, Wenhao Che, Leonidas Bleris, Chen Jia

## Abstract

In experiments, the distributions of mRNA or protein numbers in single cells are often fitted to the random telegraph model which includes synthesis and decay of mRNA or protein, and switching of the gene between active and inactive states. While commonly used, this model does not describe how fluctuations are influenced by crucial biological mechanisms such as feedback regulation, non-exponential gene inactivation durations, and multiple gene activation pathways. Here we investigate the dynamical properties of four relatively complex gene expression models by fitting their steady-state mRNA or protein number distributions to the simple telegraph model. We show that despite the underlying complex biological mechanisms, the telegraph model with three effective parameters can accurately capture the steady-state gene product distributions, as well as the conditional distributions in the active gene state, of the complex models. Some effective parameters are reliable and can reflect realistic dynamic behaviors of the complex models, while others may deviate significantly from their real values in the complex models. The effective parameters can also be applied to characterize the capability for a complex model to exhibit multimodality. Using additional information such as single-cell data at multiple time points, we provide an effective method of distinguishing the complex models from the telegraph model. Furthermore, using measurements under varying experimental conditions, we show that fitting the mRNA or protein number distributions to the telegraph model may even reveal the underlying gene regulation mechanisms of the complex models. The effectiveness of these methods is confirmed by analysis of single-cell data for *E. coli* and mammalian cells. All these results are robust with respect to cooperative transcriptional regulation and extrinsic noise. In particular, we find that faster relaxation speed to the steady state results in more precise parameter inference under large extrinsic noise.

## Introduction

Recent experiments have revealed a large cell-to-cell variation in the numbers of mRNA and protein molecules in isogenic populations due to stochasticity in gene expression and the low copy numbers of DNA and important regulatory molecules [1–3]. Live-cell imaging approaches allow a direct visualization of stochastic bursts of gene expression in living cells [4]. However these experiments are challenging and more commonly one measures the mRNA or protein expression in individual cells using flow cytometry, single-molecule fluorescence *in situ* hybridization (smFISH) [4], and single-cell RNA sequencing (scRNA-seq) [5]. Together with mathematical models, large amounts of single-cell gene expression data have been used to understand stochastic gene regulation in various biological problems, ranging from genetic engineering [6, 7] to cell fate decision [8, 9] and therapeutic targets of disease [10, 11].

Experimentally, the distributions of mRNA and protein numbers are often fitted to the predictions of mathematical models [12–18]. The most common and well-studied model of this type is the random telegraph model [19–21], which is composed of four effective reactions (Fig. 1(a))

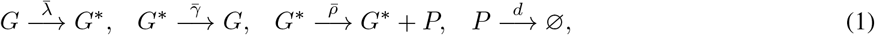

where the first two reactions describe switching of the gene between an active state *G*^*∗*^ and an inactive state *G*, the third reaction describes synthesis of the gene product *P*, that can be either mRNA [12] or protein [20], when the gene is active, and the fourth reaction describes decay of the gene product either due to active degradation or due to dilution during cell division [22, 23]. The chemical master equation describing the two-state telegraph model can be exactly solved in steady-state and in time [20, 24]. Extensions of this model to include more than two gene states have also been considered [25–27]. With tremendous efforts of quantitative and qualitative analysis, the telegraph model has been successfully applied to understand stochasticity in gene expression through (i) clarifying the biological origins of different distribution shapes [28], (ii) performing a fast and reliable inference of all parameters [17], and (iii) unravelling the gene regulation mechanisms in response to environmental changes [29].

**Figure 1:**
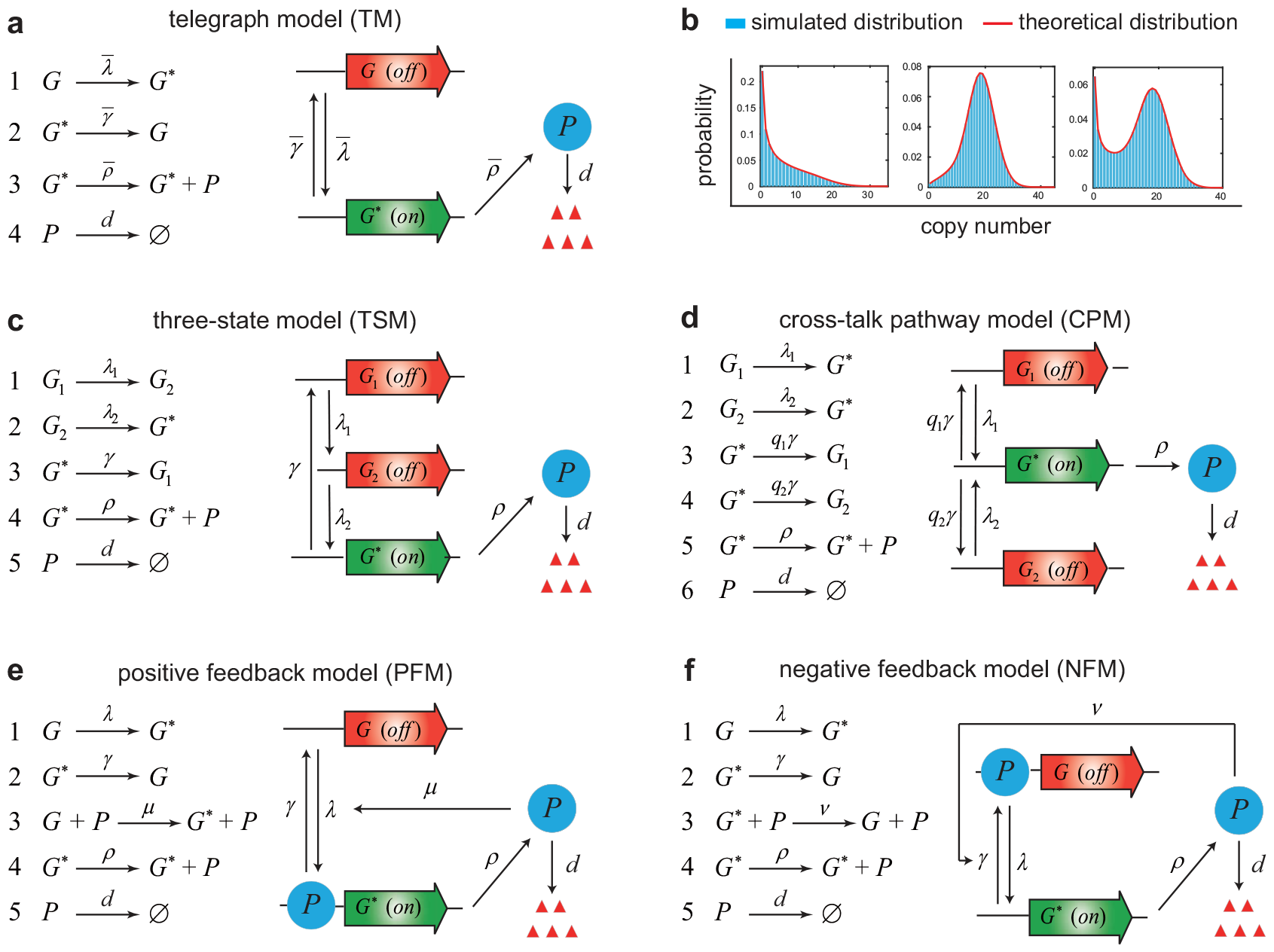
The simple telegraph model and four relatively complex gene expression models. **(a)** In the telegraph model (TM), the gene switches between an inactive (off) and an active (on) state with rates 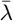 and 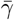. The gene product (mRNA or protein, denoted by *P* ) is synthesized with rate 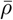 when the gene is active, and is degraded with rate *d*. **(b)** The telegraph model can generate three different shapes of steady-state distributions: a unimodal distribution with a zero peak (left panel), a unimodal distribution with a nonzero peak (middle panel), and a bimodal distribution with both a zero and a nonzero peak (right panel). **(c)** In the three-state model (TSM), the gene exhibits a “refractory” behavior: after leaving the active state with rate *γ*, the gene has to progress through two sequential inactive states with rates *λ*_1_ and *λ*_2_ before becoming active again. **(d)** In the cross-talk pathway model (CPM), the gene can be activated via two signalling pathways with rates *λ*_1_ and *λ*_2_. The competition between the two pathways is modelled by equipping them with two selection probabilities *q*_1_ and *q*_2_ = 1*−q*_1_. **(e)** In the positive feedback model (PFM), the protein produced from the gene activates its own expression with feedback strength *μ*. **(f)** In the negative feedback model (NFM), the protein produced from the gene inhibits its own expression with feedback strength *ν*.

In experiments, there are three commonly observed patterns for the mRNA or protein distributions: a unimodal distribution with a zero peak, a unimodal distribution with a nonzero peak, and a bimodal distribution with both a zero and a nonzero peak (Fig. 1(b)) [28]. Actually, the telegraph model can only produce the above three shapes of distributions. Among these three shapes, the bimodal distribution attracts the most attention since it separates isogenic cells into two distinct phenotypes [30, 31]. Bimodality of mRNA or protein distributions has been used to describe the bet-hedging strategy in microorganisms [32, 33], and to quantify cell fate decisions such as the differentiation of embryonic stem cells [34] and the activation of HIV latency [11]. For the telegraph model, it has been shown that the occurrence of bimodality requires relatively slow rates of gene state switching — a bimodal distribution can only occur when both the gene activation and inactivation rates are smaller than the decay rate [35].

In previous studies, the mRNA or protein distributions for various genes (and even for the whole genome) are often fitted to the telegraph model [12–18], by which one can obtain estimates of the rates of the underlying gene expression processes. One of the most prevalent methods of parameter inference is the maximum likelihood method which maximizes the log-likelihood function [16, 17]

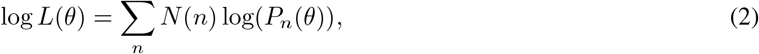

where *θ* is the parameter set, *N* (*n*) denotes the number of cells with *n* copies of mRNA or protein, and *P*_*n*_(*θ*) denotes the mRNA or protein distribution with parameter set *θ*. The decay rate *d* can be determined by measuring the half-life of mRNA or protein and the cell cycle duration [36]. However, it rarely measured in experiments and hence what is often estimated are the parameters 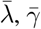, and 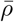 normalized by *d* [17]. To achieve fast and accurate estimation of 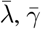, and 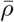, one important step is to select their initial values 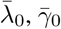, and 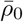 for optimization. A common choice is to set 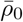 to be the maximum number of mRNA or protein molecules among single cells [16]. Once 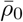 is determined, the initial values of the other two parameters, 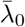 and 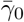, can be determined by matching the mean and variance of gene product fluctuations [16, 18].

By fitting gene expression data to the telegraph model, one can understand how all parameters change in response to varying experimental conditions [12, 15, 37]. Previous studies have revealed rich gene regulation mechanisms under different induction conditions or promoter architectures. For instance, the up-regulation of gene expression levels can be achieved by increasing the gene activation rate 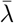 for zinc-induced yeast ZRT1 gene [6], decreasing the gene inactivation rate 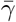 for over 20 *Escherichia coli* (*E. coli*) promoters under different growth conditions [7, 13], increasing the synthesis rate 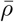 for serum-induced mammalian *ctgf* gene [38], or a combined effect of both the burst frequency 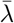 and burst size 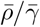 in prokaryotic and eukaryotic cells [29, 37, 39].

However, the conventional telegraph model is limited in its predictive power because it lacks a description of some important biological mechanisms such as feedback regulation, non-exponential gene inactivation durations, and multiple gene activation pathways (Fig. 1(c)-(f)). The telegraph model can only be used to study genes that are unregulated, and it fails for regulated genes. One of the most common gene network motifs is an autoregulatory feedback loop whereby protein expressed from a gene activates or represses its own transcription (Fig. 1(e),(f)) [40–43]. It has been estimated that 40% of all transcription factors self-regulate in *E. coli* [44] with most of them participating in negative autoregulation [45]. An effective method of inferring the sign of autoregulation has been proposed based on gene expression measurements under different feedback strengths [46].

Except feedback regulation, another important mechanism that regulates gene expression is non-exponential gene inactivation periods. In the telegraph model, the time spent in the active or inactive gene state has an exponential distribution. The exponential active period is generally a reasonable assumption [47]. However, recent studies in mammalian and bacterial cells have shown that the inactive periods for some genes may have a non-exponential peaked distribution [48–51]. This suggests that the gene dynamics in the inactive period may contain two rate-limiting steps and exhibit a “refractory” behavior: after leaving the active state, the promoter has to progress through two inactive states before becoming active again (Fig. 1(c)). This refractory behavior is probably due to the fact that the activation of the promoter is a complex multi-step biochemical process due to chromatin remodeling and the binding and release of transcription factors or RNA polymerase [52]. Different Bayesian methods have been applied to estimate all parameters of this refractory model based on time-course gene expression data [53, 54].

Another possible mechanism that regulates gene expression is the existence of multiple signalling pathways during gene activation [47, 55]. In the telegraph model, there is only one gene activation pathway. Recent studies [56, 57] have shown that the competition between two gene activation pathways (Fig. 1(d)) can well capture the rapid overshooting behavior of transcription levels observed in mouse fibroblasts under the induction of tumor necrosis factor [58]. Such behavior cannot be explained by a single gene activation pathway with one or more rate-limiting steps since it either generates monotonic transcription dynamics or triggers a long lag to reach the peak of the transcription level. Moreover, the existence of two gene activation pathways can also capture the time-course mRNA expression data observed for yeast *HSP12* gene under NaCl osmotic stress which exhibit unimodal distributions with a zero peak for small and large times, while exhibit bimodal distributions for intermediate times [59, 60]. Such dynamic transitions between different distribution shapes are rarely observed in the telegraph model and other gene expression models [61–63].

Integrating the above biological mechanisms into the telegraph model can generate more complex models of stochastic gene expression (Fig. 1(c)-(f)). An essential problem is, compared to the telegraph model, there is still a lack of effective methods of theoretical analysis and parameter inference for these models due to the increased complexity of model structures and increased number of model parameters. Furthermore, it is also difficult to distinguish these relatively complex models from the simple telegraph model since they often exhibit similar distribution shapes. This raises the questions of (i) whether some parameters for a complex model can be accurately inferred, (ii) whether we can distinguish a complex model from the telegraph model, and (iii) whether we can infer the gene regulation mechanism of a complex model by using gene expression data under different induction conditions.

In this paper, we will provide insights to these questions. Our strategy is not to investigate the complex models themselves; rather, we examine these models by fitting their steady-state distributions to the telegraph model and then obtain estimates of the “effective” parameters. In fact, the idea of fitting a complex model to the telegraph model has been previously carried out in [17], where the authors realized that the three-state model shown in Fig. 1(c) may be more accurate in mammalian cells but they still fitted the mRNA distributions for thousands of genes to the telegraph model since, as explained in [17], “the resulting steady-state distribution for the extended (three-state) model is very close to the two-state model and to distinguish between these similar models, additional information such as multiple time measurements within the same cell is needed.” In general, the estimated values of the parameters (the mRNA or protein synthesis rate and the gene activation and inactivation rates) in the “artificial” telegraph model may deviate largely from their real values in the complex models. However, we find that the effective parameters are still sometimes reliable and can also reveal important dynamical properties of the complex models such as the ability for a complex model to produce bimodality. Furthermore, using additional information such as gene expression data at multiple time points or measurements under varying experimental conditions, we provide an effective method of distinguishing the complex models from the telegraph model, and we also show that fitting the mRNA or protein distributions to the telegraph model may even reveal the underlying gene regulation mechanisms of the complex models. The effectiveness of these methods is confirmed by analysis of published data for *E. coli* and mammalian cells. All these results are shown to be robust with respect to cooperative transcriptional regulation and extrinsic noise.

## Results

### Four relatively complex gene expression models revisited

Here we recall four relatively complex models of stochastic gene expression including the three-state model, cross-talk pathway model, positive feedback model, and negative feedback model (see Fig. 1(c)-(f) for illustration and the detailed reaction schemes). All of them have been extensively studied in the literature and are established by integrating a particular genetic regulation mechanism into the telegraph model. Like the telegraph model, all the four complex models are composed of gene state switching, synthesis of the gene product, and decay of the gene product either due to active degradation and dilution during cell division. The gene product for the former two models can be either mRNA [12] or protein [20], while for the latter two models, the gene product must be protein since feedback regulation is realized by binding of proteins to the promoter.

The three-state model assumes that the process of gene activation contains two rate-limiting steps (Fig. 1(c)); this explains the non-exponential gene inactivation period observed in experiments [48–51]. The dynamics of the three-state model is controlled by two consecutive gene activation steps with rates *λ*_1_ and *λ*_2_, gene inactivation rate *γ*, synthesis rate *ρ* of mRNA or protein, and decay rate *d*. For convenience, we set *d* = 1 in what follows. This is not an arbitrary choice but stems from the fact that the time and parameters can always be non-dimensionalized using *d*. Specifically, the time given below should be understood to be non-dimensional and equal to the real time multiplied by *d*, while the parameters *λ*_*i*_, *γ*, and *ρ* given below should also be understood to be non-dimensional and equal to their real values divided by *d*.

The cross-talk pathway model describes competitive binding of two transcriptional factors to the promoter: one actives the gene via a weak signalling pathway with rate *λ*_1_, and the other actives the gene via a strong pathway with a larger rate *λ*_2_ *> λ*_1_ (Fig. 1(d)) [64, 65]. The competition between the two pathways is modelled by equipping them with two selection probabilities *q*_1_ and *q*_2_ satisfying *q*_1_ + *q*_2_ = 1. In other words, the gene is activated via the weak pathway with probability *q*_1_ and is activated via the strong pathway with probability *q*_2_. The mRNA or protein is synthesized with rate *ρ* and is degraded with rate *d* = 1. This model has been successfully used to explain the rich transcription dynamics observed in mouse fibroblasts and yeast under different induction conditions [57, 60].

One of the most common gene network motifs is an autoregulatory feedback loop whereby protein produced from a gene activates or represses its own expression [45]. It has been estimated that 40% of all transcription factors self-regulate in *E. coli* [44]. The positive feedback model describes an autoregulatory loop whereby protein expressed from a gene activates its own transcription (Fig. 1(e)). It has the same reaction scheme as the telegraph model except that the protein activates the gene with rate constant *μ*, which characterizes the strength of positive feedback. Note that it reduces to the telegraph model when *μ* = 0.

Among the 40% transcription factors that regulate their own expression in *E. coli*, most of them participate in negative autoregulation [45]. The negative feedback model describes an autoregulatory loop whereby protein expressed from a gene represses its own transcription (Fig. 1(f)). It has the same reaction scheme as the telegraph model except that the protein represses the gene with rate constant *ν*, which characterizes the strength of negative feedback. In fact, the steady-state gene product distributions for the four relatively complex models can all be solved analytically and the exact distributions can be found in Supplementary Sec. 1.

Note that there are three parameters for the telegraph model (assuming that *d* = 1), four parameters for the three-state, positive feedback, and negative feedback models, and five parameters for the cross-talk pathway model. It has been shown recently that the distributions of protein numbers in *E. coli* measured using single-molecule fluorescence microscopy often have a unimodal distribution [3] and the distributions of mRNA numbers measured using scRNA-seq often have a negative binomial or zero-inflated negative binomial distribution [66–68]. Given these (relatively simple) distributions, it is almost impossible to accurately infer all the four or five parameters of a complex model. A solution of this is to fit the experimental distributions of mRNA or protein numbers to a simple telegraph model, by which one can obtain estimates of the three “effective” parameters 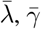, and 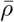 [12–17]. However, it is not clear whether a simple telegraph model can always capture the distribution of a complex model, and it is also not clear whether these effective parameters can reflect the realistic gene expression processes behind a complex model.

### The telegraph model can accurately capture the distributions of complex models

We first examine whether the gene product distribution of a complex model can be well approximated by that of the telegraph model. To this end, for each of the four complex models, we generate synthetic data of mRNA or protein numbers for *N* = 10^4^ cells using the stochastic simulation algorithm (SSA), and then fit the steady-state simulated distribution to the telegraph model using the maximum-likelihood method [12–17] (Fig. 2(a)). The detailed description of the method can be found in Supplmentary Sec. 2. Here the gene activation rate of the weak signalling pathway for the cross-talk pathway model is fixed to be *λ*_1_ = 0.2 so that each complex model has four independent parameters. To proceed, we proportionally select five different values for each of the four parameters, which cover large swathes of parameter space and give 5^4^ = 625 different parameter sets for each complex model (see Methods).

**Figure 2:**
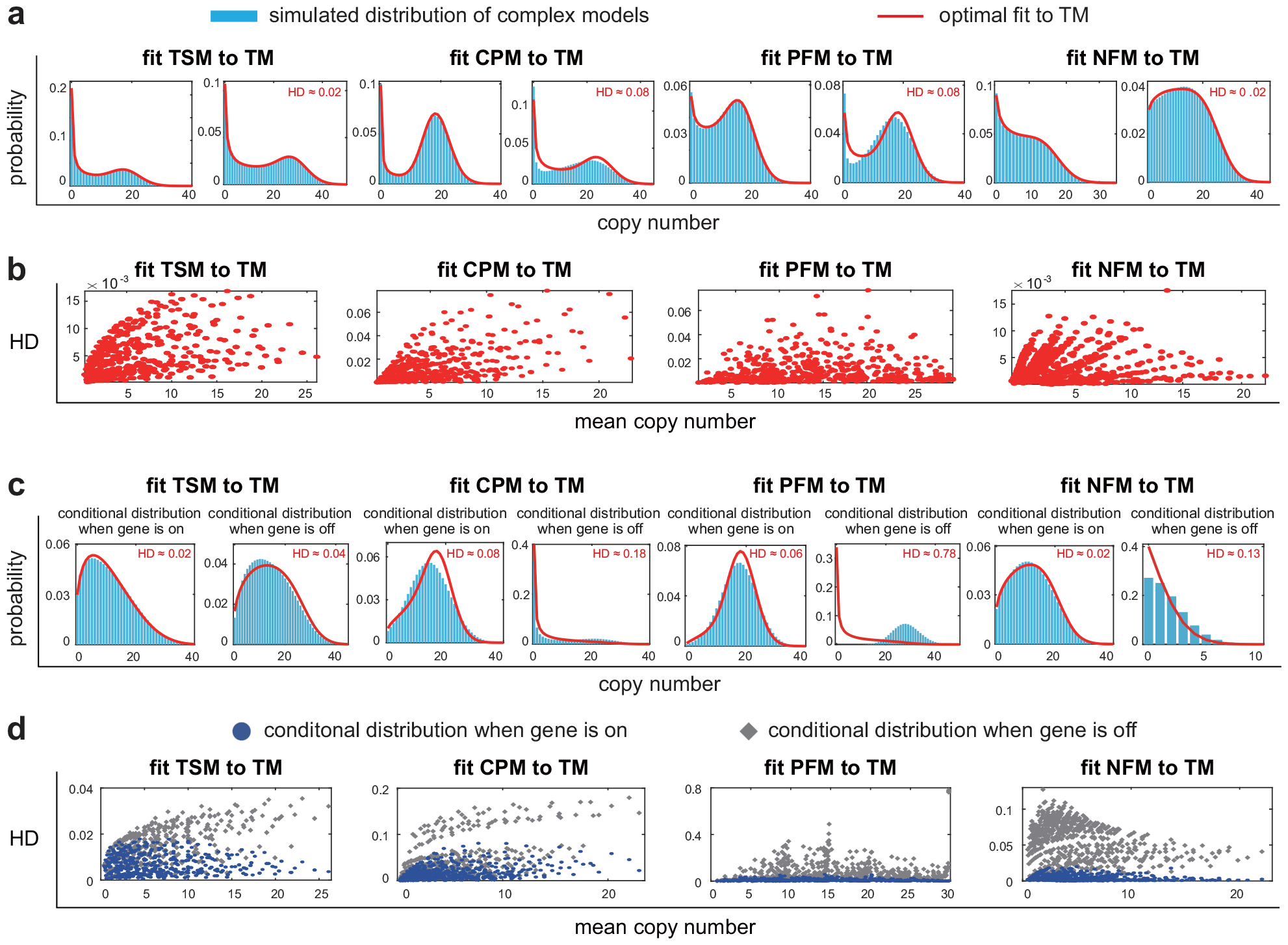
Fitting the steady-state distributions of complex models to the simple telegraph model. For each complex model, synthetic data of gene product numbers are generated using the SSA under 625 parameter sets. **(a)** In steady state, all the simulated distributions (blue bars) are well captured by the predictions of the effective telegraph model (red curve). For each complex model, the left panel shows a typical gene product distribution and the right panel shows the distribution with worse telegraph model approximation, i.e. maximum HD value. **(b)** For each complex model, the HD between the simulated distribution and its telegraph model approximation is shown as a function of the mean expression level for the 625 parameter sets. The HD is less than 0.08 for all complex models. **(c)** In steady state, the telegraph model not only captures the total gene product distribution of a complex model, but also captures the conditional distribution in the active gene state. In contrast, for all complex models except the three-state model, the conditional distribution in the inactive gene state in general fails to be captured by the telegraph model. For each complex model, the left (right) panel shows the conditional distribution when the gene is on (off) with worse telegraph model approximation, i.e. maximum HD value. **(d)** For each complex model, the HD is shown as a function of the mean expression level for the 625 parameter sets. The blue circles (grey diamonds) show the HD between the conditional distribution when the gene is on (off) and its telegraph model approximation. The maximum HD for blue circles is only 0.08 for all complex models, while the maximum HD for grey diamonds can be as large as 0.78.

Like the telegraph model, each complex model can generate unimodal or bimodal distributions of gene product numbers (Fig. 2(a)). For each of the 625 parameter sets, the synthetic data obtained using the SSA are then fitted to the telegraph model. Interestingly, we find that the simulated distributions for all complex models and all parameter sets can be well approximated by the predictions of the telegraph model with the Hellinger distance (HD) between the two distributions always less than 0.08 (Fig. 2(a),(b)) and with the Kullback-Leiber divergence (KLD) between the two distributions always less than 0.025 (Supplementary Fig. S1). This shows that the distributions of complex models can generally be well captured by the telegraph model with effective parameters 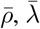, and 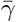. In what follows, the telegraph model equipped with the effective parameters is referred to as the *effective telegraph model* of a complex model. Intriguingly, both the HD and KLD seem to positively correlate with mean gene product number (Fig. 2(b)). A possible reason is that a low mean copy number is usually associated with a unimodal distribution with a zero peak that is easy to be captured by the telegraph model, while a high mean copy number usually corresponds to a distribution with a nonzero peak that is more difficult to be captured by the telegraph model, leading to a higher HD or KLD.

While the steady-state distributions of complex models can be well fitted by the effective telegraph model, it is not clear whether the conditional distributions of complex models in the inactive and active gene states can also be captured by the effective telegraph model. Specifically, let *P*_*i,n*_ denote the steady-state probability of observing *n* copies of the gene product when the gene is in state *i*, with *i* = 0, 1 corresponding to the inactive and active states, respectively. Then the conditional gene product distribution in gene state *i* can be calculated as

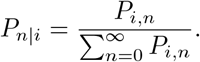

For each complex model, we generate the conditional distributions in the inactive and active gene states (*P*_*n*|0_ and *P*_*n*|1_) using the SSA under all 625 parameter sets. Similarly to the total gene product distribution (*P*_*n*_ = *P*_0,*n*_ + *P*_1,*n*_), the conditional distribution in the active gene state (*P*_*n*|1_) for each complex model can always be well approximated by the predictions of the effective telegraph model with an HD less than 0.08 (Fig. 2(c) and the blue dots in Fig. 2(d)). The situation is different for the inactive gene state. We find that for each complex model except the three-state model, the conditional distribution in the inactive gene state (*P*_*n*|0_) fails to be captured by the effective telegraph model, manifested by significantly larger HD values. The worst approximation occurs for the positive feedback model where the HD can be as large as 0.78 (Fig. 2(c) and the grey dots in Fig. 2(d)). According to our simulations, poor approximations generally occur when the gene is mostly in the active state. To explain this, note that the total gene product distribution can be represented as 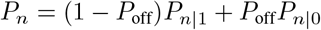, where 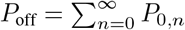 is the probability of the gene being in the inactive state. When gene is mostly on, we have *P*_off_ « 1. In this case, even if the effective telegraph model can capture both *P*_*n*_ and *P*_*n*|1_, it fails to capture *P*_*n*|0_ because *P*_off_ is too small.

### Linking effective parameters to realistic gene expression processes

In the telegraph model, 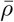 represents the synthesis rate of the gene product, while 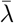 and 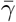 represent the frequencies of the gene being activated and inactivated, respectively (Fig. 1(a)). In other words, 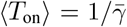 and 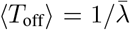 represent the mean active and inactive durations of the gene, respectively. However, it is not clear whether the effective parameters 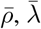, and 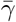 of the four complex models can reflect the same dynamic properties.

Note that the synthesis rate is *ρ* for each complex model (Fig. 1(c)-(f)). However, the mean holding times in the active and inactive states for the four complex models have completely different expressions. For the three-state model, since gene inactivation consists of only one exponential step and gene activation consists of two exponential steps, the mean active and inactive durations can be easily calculated as

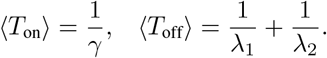

For the cross-talk pathway model, since there is only one pathway for gene inactivation and two pathways for gene activation, the mean active and inactive durations can be easily calculated as

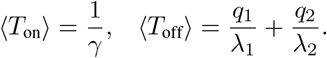

The expressions of the mean holding times for positive and negative feedback models are much more complicated and the detailed expressions can be found in Supplementary Sec. 3.

Next we examine whether the three effective parameters of a complex model can reflect the realistic gene expression processes. To this end, we consider the relative error between 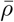 and *ρ*, the relative error between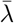 and 1*/*(*T*_off_), and the relative error between 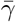 and 1*/*(*T*_on_), i.e.

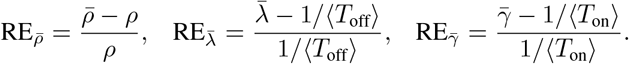

Moreover, we say that an effective parameter is *over-estimated* (*under-estimated*) if the corresponding relative error is greater (less) than zero. For each complex model, we calculate the relative errors of the three effective parameters under all 625 parameter sets which are chosen to be the same as in Fig. 2(a). Fig. 3(a) illustrates the sample mean and standard deviation of 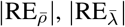, and 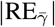 for all parameter sets (also see Supplementary Fig. S2 for the empirical distributions of 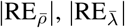, and 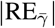). In addition, Fig. 3(a) also shows the empirical proportions of 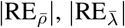, and 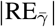 being greater than 0.2. If the relative error of an effective parameter has an absolute value less than 0.2, then we believe that it can reflect the realistic dynamic property of the corresponding complex model.

**Figure 3:**
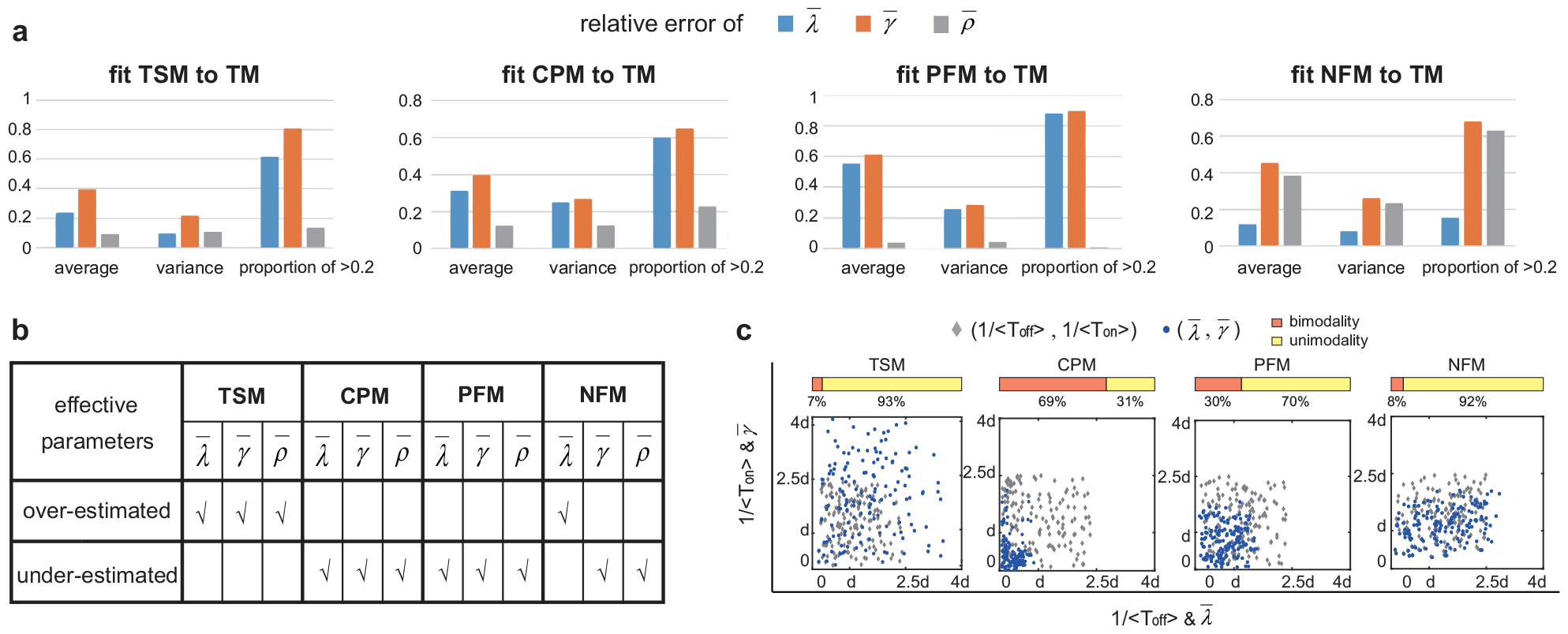
Linking effective parameters to their real values in complex models. **(a)** For each complex model, the absolute values of relative errors of the three effective parameters 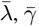, and 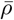 are computed under 625 parameter sets, along with their sample means, sample variances, and the sample frequencies of relative errors being greater than 0.2. The effective parameter 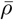 is closed to the synthesis rate *ρ* for the three-state, cross-talk pathway, and positive feedback models, while the effective parameter 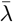 is closed to the gene activation rate *λ* for the negative feedback model. **(b)** Accuracy of the three effective parameters 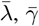, and 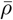 for each complex model. For the three-state model, all effective parameters are over-estimated; for the cross-talk pathway and positive feedback models, all effective parameters are under-estimated; for the negative feedback model,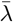 is over-estimated, while 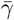 and 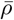 are under-estimated. **(c)** For each complex model, 150 parameter sets are randomly generated such that 1*/ ⟨T*_off_*⟩* and 1*/ ⟨T*_on_*⟩* are between 0 and 2.5*d* (grey diamonds). For the three-state model, the scatter plot of 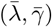 escapes from the potential bimodal region of 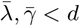; for the cross-talk pathway and positive feedback models, the scatter plot of 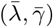 moves towards the potential bimodal region; for the negative feedback model, the scatter plot of 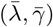 neither escapes from nor moves towards the potential bimodal region. The yellow (orange) bar shows the proportion of parameter sets that give rise to a unimodal (bimodal) distribution.

For the three-state model, the sample mean of 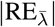 and 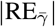 are large, while 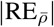 has a relatively small sample mean; in particular, there are only 13.6% of parameter sets such that 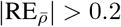. This suggests that in most cases, the estimated value of 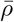 is very close to the real synthesis rate *ρ*. Similar phenomenon is also observed for the cross-talk pathway and positive feedback models. In particular, for the positive feedback model, almost all values of 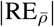 are less than 0.2, suggesting that fitting the steady-state protein distribution of the positive feedback model to the telegraph model can always provide a reliable estimation of the synthesis rate *ρ*. The situation is different for the negative feedback model, where both 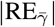 and 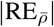 have a relatively large sample mean, while 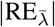 has a relatively small sample mean. This suggests that the estimated value of 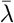 can reflect the realistic gene activation rate of the negative feedback model. Interestingly, for all complex models, the gene inactivation frequency is the worst estimated parameter when fitted to the telegraph model. This is consistent with the results obtained in [69], which makes an extensive investigation of the accuracy of parameter estimation using the telegraph model.

Interestingly, our simulations also reveal that the relative errors of the three effective parameters follow some consistent principles: (i) for the three-state model, all effective parameters are over-estimated; (ii) for the cross-talk pathway and positive feedback models, all effective parameters are under-estimated; (iii) for the negative feedback model, 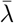 is over-estimated, while 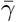 and 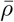 are under-estimated. Here the principles are consistent in the sense that it is impossible that an effective parameter is over-estimated for a certain parameter set, while it is under-estimated for another parameter set. These rules of over-estimation and under-estimation are summarized in Fig. 3(b). These rules can be further used to characterize the ability for a complex model to exhibit bimodality. For each complex model, we randomly select 150 parameter sets such that the values of 1*/⟨T*_off_*⟩* and 1*/⟨T*_on_*⟩* are between 0 and 2.5*d* (see Methods), and for each parameter set, we fit the synthetic data obtained from the SSA to the telegraph model. The values of the real gene switching frequencies 1*/⟨T*_off_*⟩* and 1*/⟨T*_on_*⟩* are shown by the grey dots in Fig. 3(c). We then superpose the values of the effective parameters 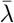 and 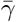 for the 150 parameter sets (shown by the blue dots) onto the same figure.

For the telegraph model, it has been proved that a bimodal distribution can only occur when both gene switching rates are smaller than the decay rate, i.e. 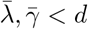 [35]. The principles summarized in Fig. 3(b) show that 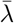 and 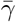 are under-estimated for the cross-talk pathway and positive feedback models. This is also shown in Fig. 3(c), where the scatter plots of 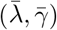 for the two models move towards the potential bimodal region, i.e. 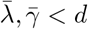. Hence compared to the telegraph model, the cross-talk pathway and positive feedback models are more likely to exhibit bimodality. This is consistent with experimental observations [70, 71] and is possibly due to the fact that cross-talk pathway and positive feedback tend to increase gene expression noise [46]. In contrast, both 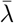 and 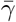 are over-estimated for the three-state model (Fig. 3(b)) and thus the scatter plot of 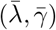 tends to escape from the potential bimodal region (Fig. 3(c)). This shows that the three-state model is less likely to display bimodality, possible due to the fact that a multi-step gene activation process reduces gene expression noise [72]. For the negative feedback model, 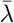 is over-estimated and 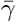 is under-estimated (Fig. 3(b)). Hence the scatter plot of 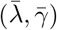 neither moves towards nor escape from the potential bimodal region (Fig. 3(c)). This indicates that negative autoregulation has weak influence on bimodality. Fig. 3(c) also shows the proportion of parameter sets that lead to a unimodal or bimodal distribution for each complex model. Again, the fraction of bimodal distributions is significantly higher for the cross-talk pathway and positive feedback models.

Here we show that the ability for a complex model to produce bimodality is closely related to the under-estimation of the effective gene activation and inactivation rates, 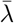 and 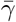, when fitted to the telegraph model. This is consistent with previous findings that slow gene state switching is an important source of bimodality [40, 73] and has the potential to be used to analyze bimodality for more complex gene expression models.

### Identification of complex models using snapshot data at multiple time points

Given the experimental distributions of mRNA or protein numbers in steady state, it is almost impossible to distinguish a complex model from the telegraph model since the latter can accurately capture the steady-state distribution of the former. To further test this, for each complex model, we generate synthetic data of mRNA or protein numbers using the SSA for *N* = 10^2^, 10^3^, 10^4^, 10^5^ cells under 625 parameter sets. Then we fit the steady-state distribution obtained from the SSA to the complex model and the telegraph model, respectively, by maximizing the log-likelihood function *L*(*θ*). To distinguish between the two competing models, a common strategy is to select the model with lower corrected Akaike information criterion (AICc) [74]

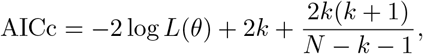

where *k* is the number of parameters (*k* = 4 for each complex model and *k* = 3 for the telegraph model) and *N* is the sample size. Here we use the AICc because it imposes greater penalty on the number of parameters than the conventional AIC, especially when the sample size is small. According to simulations, the proportion of incorrect model selection, i.e. the telegraph model has a smaller AICc, decreases with the sample size *N* . For each complex model, over 90% of parameter sets lead to incorrect model selection for *N* = 10^2^ cells (typical sample size for smFISH and scRNA-seq data), and the proportion is still over 40% even for *N* = 10^4^ cells (Supplementary Fig. S3). This clearly shows that reliable model selection fails to be made based solely on steady-state data.

To distinguish a complex model from the telegraph model, additional information such as snapshot data at multiple discrete time points within the same cell population measured using e.g., live-cell imaging, flow cytometry, smFISH, and scRNA-seq, is needed [75, 76]. Recent studies have proposed various statistical methods, such as the maximum likelihood method [59, 75] and various Bayesian method [53, 54, 77], to search optimal kinetic parameters based on single-cell data at multiple time points. However, no matter which method is used, the first and most important step is to determine which model (the telegraph model or more complex models) is the most competitive to describe the snapshot data [59].

We next examine how to distinguish a complex model from the telegraph model by using snapshot data. Here we assume that initially there is no gene product molecules in the cell and the gene is in the inactive state. This mimics the situation where the gene has been silenced by some repressor over a period of time such that all gene product molecules have been removed via degradation. At time *t* = 0, the repressor is removed and we investigate how gene expression recovers. Note that a complex model and its effective telegraph model have very similar steady-state distributions; however, they may exhibit completely different dynamic behaviors since their time-dependent distributions are generally different. For each complex model, we compute the time-dependent mean *M* (*t*) and variance *σ*^2^(*t*) of gene product fluctuations using the finite-state projection (FSP) algorithm [75] under all 625 parameter sets which are chosen to be the same as in Fig. 2. The time-dependent mean and variance for the effective telegraph model are denoted by 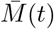 and 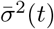, respectively.

Interestingly, for the three-state and positive feedback models, we find that the mean curve *M* (*t*), as a function of time *t*, is always below its counterpart 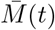 for the effective telegraph model for all parameter sets. In contrast, the mean curve *M* (*t*) for the cross-talk pathway and negative feedback models is always above its counterpart 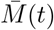 for the effective telegraph model (Fig. 4(a)). This is probably because compared to the telegraph model, the three-state and positive feedback models have slower relaxation speed to the steady state, while the cross-talk pathway and negative feedback models relax to the steady state faster [44, 78]. In particular, the cross-talk pathway model may even perform overshooting behavior [79], where the maximum value of the mean curve *M* (*t*) exceeds its steady-state value (blue dashed curve in Fig. 4(a)). Similar dynamic features are also observed for the time-dependent second moment ⟨*n*^2^⟩ (*t*); however, common indicators of gene expression noise, such as the coefficient of variation squared *σ*^2^(*t*)*/M* ^2^(*t*) and the Fano factor *σ*^2^(*t*)*/M* (*t*), present less easily distinguishable dynamic differences between a complex model and its effective telegraph model (Supplementary Fig. S4).

**Figure 4:**
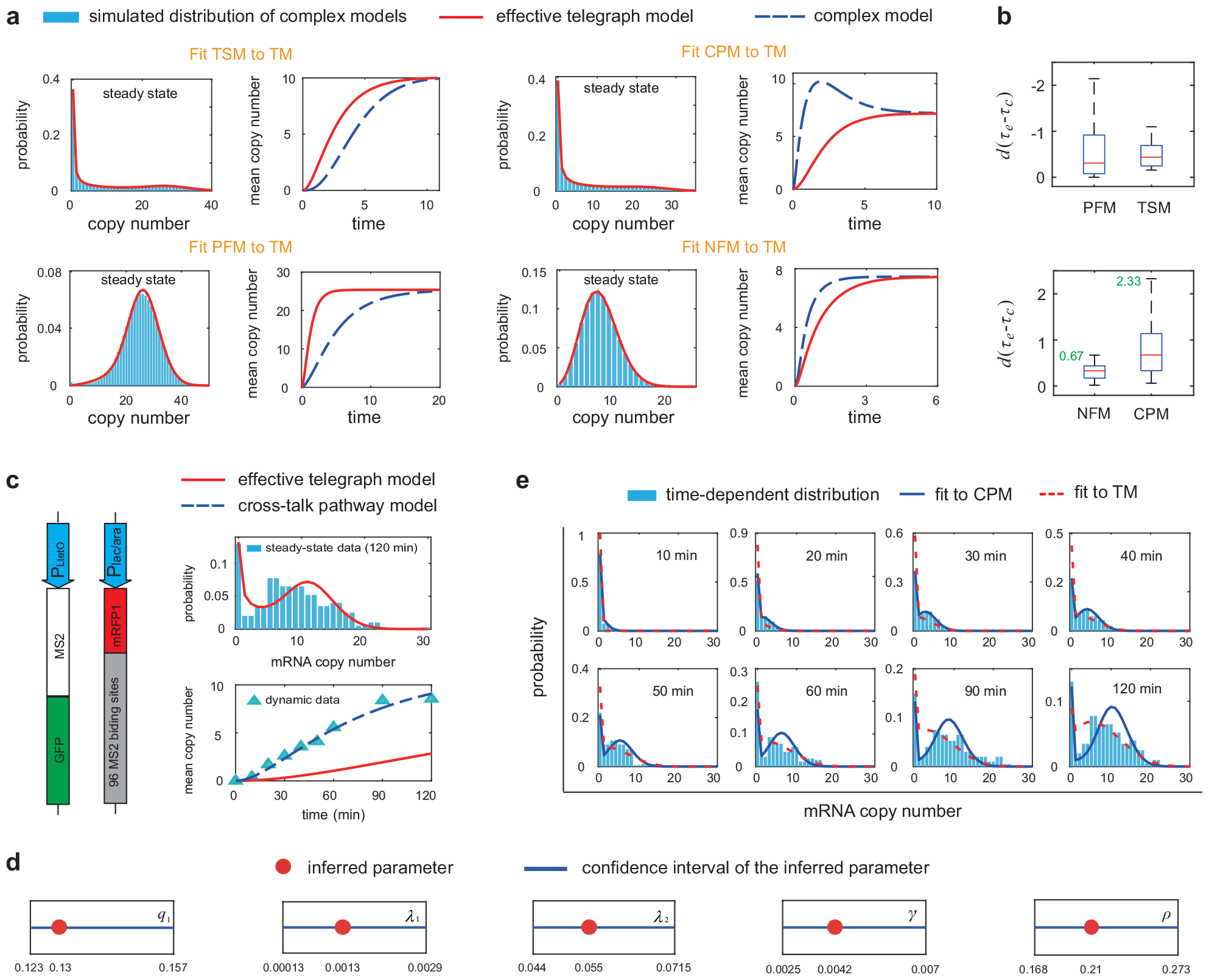
Determining the most competitive model to describe single-cell data at multiple time points. **(a)** The three-state and positive feedback models have smaller time-dependent mean curve compared to the effective telegraph model, while the cross-talk pathway and negative feedback models have larger time-dependent mean curve. **(b)** Box plots of *d*(*τe−τ_c_*) for each complex model, where the *τ*_*c*_ is the response time for a complex model and *τ*_*e*_ is the response time for the effective telegraph model. Here the response time is defined as the time for the mean curve to reach half of its steady-state value [44]. **(c)** In *E. coli* cells, the mRNA of interest, under the control of an inducible promoter P_lac*/*ara_, was consisted of the coding region for a red fluorescent protein mRFP1, followed by a tandem array of 96 MS2 binding sites (left panel) [1]. The GFP, independently produced from the promoter P_LtetO_, tagged the target transcript by binding to the MS2 binding sites. The number of target transcripts in a single cell was computed using fluorescence intensities of GFP at nine time points from 0 - 120 min. The steady-state mRNA distribution (at 120 min) was fitted to the telegraph model with measured decay rate *d* = 0.014 min^*−*1^ [1] (upper-right panel) and the three effective parameters are estimated to be 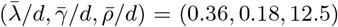. The real time-dependent mean expression levels (blue triangles in the lower-right panel) are much larger than the mean expression levels predicted by the effective telegraph model (red curve), suggesting that the cross-talk pathway model is a potential candidate to describe the data. **(d)** Point estimates (red points) and confidence intervals (blue lines) for the six parameters *q*_1_, *q*_2_, *λ*_1_, *λ*_2_, *γ*, and *ρ* when fitting the data to the cross-talk pathway model. Here the confidence intervals are computed using the profile likelihood method. **(e)** The cross-talk pathway model (blue curves) provides a much better fit of the time-dependent mRNA distributions than the telegraph model (red dashed curves). The parameters for the cross-talk pathway model are estimated to be *q*_1_ = 0.13, *q*_2_ = 1 *− q*_1_, and (*λ*_1_, *λ*_2_, *γ, ρ*) = (0.0013, 0.055, 0.0042, 0.21) min^*−*1^.

The above results provide an effective method of distinguishing a complex model from the telegraph model. Given single-cell data at multiple time points, we can first fit the steady-state distribution to the telegraph model and estimate the effective parameters 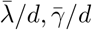, and 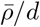, where the decay rate *d* = (log 2)*/T* + (log 2)*/T*_*c*_ can be determined by measuring the half-life *T* of the gene product and the cell cycle duration *T*_*c*_ [36]. We then compare the experimental mean curve *M* (*t*) obtained from the snapshot data and the mean curve 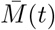 of the effective telegraph model. If the former is above the latter, then we have good reasons to believe that the cross-talk pathway or negative feedback model is more competitive. In contrast, if the former is below the latter, then we may select the three-state or positive feedback model to describe the data.

To gain deeper insights, for each parameter set, we compute the response time *τ*_*c*_ for a complex model and the response time *τ*_*e*_ for its effective telegraph model, where the response time is defined as the time required for the mean curve (*M* (*t*) or 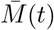) to reach half of its steady-state value [44]. From Fig. 4(a), it is clear that *τ*_*e*_ *< τ*_*c*_ for the three-state and positive feedback models and *τ*_*e*_ *> τ*_*c*_ for the cross-talk pathway and negative feedback models. We find that the response time gap *G* = *d*(*τ*_*e*_ − *τ*_*c*_) plays a vital role; here we multiply the true response time gap *τ*_*e*_ − *τ*_*c*_ by *d* since we want to transform it into a non-dimensional quantity. The 625 parameter sets yield 625 values of *G*. Fig. 4(b) shows the box plots of *G* for all complex models. Interestingly, in the average sense, the cross-talk pathway model has a much larger *G* compared to the negative feedback model. This suggests that the response time gap serves as an effective indicator to distinguish between the two complex models. Note that the maximum of *G* is only 0.67 for the negative feedback model. Hence if the experimental value of *G* is larger than 0.67 (here *τ*_*c*_ should be understood as the response time obtained from the experimental mean curve), then the negative feedback model can be safely excluded.

### Validation of theoretical results using snapshot data at multiple time points

To validate our method, we apply it to the data set of mRNA expression at multiple discrete time points measured in living *E. coli* cells [1]. In this experiment, anhydrotetracycline was first added to a growing culture, which induced the P_LtetO_ promoter to produce MS2 protein fused to green fluorescent protein (GFP) (Fig. 4(c)). The mRNA target, under the control of another inducible promoter P_lac*/*ara_, was consisted of the coding region for a red fluorescent protein mRFP1, followed by a tandem array of 96 MS2 binding sites. MS2-GFP fusion protein produced from the P_LtetO_ promoter can then bind to the MS2 binding sites and hence the synthesized transcripts from the P_lac*/*ara_ promoter were tagged by GFP. In other words, the abundances of mRFP1 protein were measured by red fluorescence and the corresponding mRNA abundances were counted by green foci. The P_lac*/*ara_ promoter can be repressed by LacI and can be activated by AraC. Activation of the promoter was induced by adding arabinose to obtain full activation of the *ara* system followed by adding isopropylthio-*β*-D-galactoside (IPTG) to repress the *lac* component. Samples were imaged using fluorescence microscopy at nine different time points from 0 - 120 min, and the number of transcripts in individual cells was computed according to green foci. In what follows, we only focus on the dynamics of mRFP1 transcripts and do not consider the expression of mRFP1 protein.

All cells contained no green foci at *t* = 0 min, suggesting that initially there are no mRNA molecules. Under the induction of arabinose and IPTG, the mean number of transcripts increases monotonically and approaches the steady state at *t* = 120 min (Fig. 4(c)). The mRNA expression in steady state exhibits an apparent bimodal distribution with a zero and a nonzero peak. We then fit the steady-state mRNA distribution to the telegraph model (Fig. 4(c)) and the three effective parameters are estimated to be 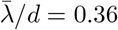, 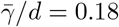, and 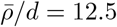. The mRNA tagged by GFP is very stable and its decay rate is measured to be *d* = 0.014 min^*−*1^ [1]. Fig. 4(c) compares the experimental mean curve *M* (*t*) and the mean curve 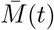 of the effective telegraph model computed using the effective parameters. It is clear that 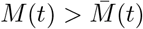 for all time points. Our theory then suggests that the cross-talk pathway and negative feedback models are more reasonable than the telegraph model. Since there is no evidence that mRFP1 protein binds to its own promoter to form an autoregulatory loop [1], we have good reasons to believe that the cross-talk pathway model may be the most competitive to describe the snapshot data (this fact will be confirmed using two different methods based on data analysis; see below).

We next estimate all the parameters of the cross-talk pathway model based on the time-dependent distributions of transcript numbers. An accurate parameter inference can be achieved by maximizing the following log-likelihood function summed over all nine time points [75]:

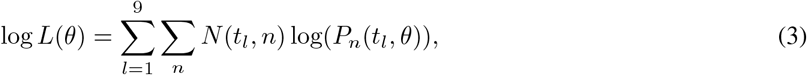

where *θ* is the parameter set for the cross-talk pathway model, *N* (*t*_*l*_, *n*) denotes the number of cells with *n* transcripts at time *t* = *t*_*l*_, and *P*_*n*_(*t*_*l*_, *θ*) denotes the theoretical mRNA distribution at time *t* = *t*_*l*_. Here the theoretical distribution is computed using FSP assuming that initially there are no mRNA molecules and the gene is off. Note that similar methods of parameter inference based on time-course measurements have been performed in [80]. To handle the positive constraint on rate parameters, we rewrite 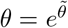 and set 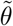 to be the optimization variables [77]. One exception is the selection probability *q*_2_ ∈ (0, 1) of the strong pathway, for which we rewrite 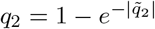 and set 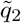 to be the optimization variable. Note that while our inference method is robust for the telegraph model (Supplementary Fig. S5), there may be ambiguity in parameter estimation for more complex models [80]. To check this, we compute the 95% confidence intervals for all parameters (Fig. 4(d)) using the profile likelihood method (see Methods). Following [81], the inference uncertainty for a given parameter is defined as the width of the confidence interval divided by the point estimate. From Fig. 4(d), the uncertainty is computed as 0.26 for *q*_1_, 2.1 for *λ*_1_, 0.5 for *λ*_2_, 1.1 for *γ*, and 0.5 for *ρ*. The parameter *λ*_1_ has a higher uncertainty than other parameters because the value of *λ*_1_ is too small so that it is difficult to precisely determine its value. These relatively low uncertainties ensure high precision of the inferred parameters.

Fig. 4(e) illustrates the experimental mRNA distributions at all measured time points and the optimal fit of these distributions to the cross-talk pathway model (blue curves) and the telegraph model (red dash curves). It can be seen that the former indeed behaves much better than the latter. First, the total HD for the cross-talk pathway model summed over all time points is 1.03, which is less than a much higher HD of 1.67 for the telegraph model. Second, the bimodal distributions after 30 min can be very well reproduced by the cross-talk pathway model but fail to be captured by the telegraph model. To reinforce our result, we also fit the time-dependent mRNA distributions to the negative feedback model. Interestingly, the estimated negative feedback strength *ν* is always zero for 50 sets of initial optimization parameters. This again shows that the negative feedback model fails to capture the time-course data since it is even worse than the telegraph model. In addition, we also compute the experimental response time *τ*_*c*_ and the response time *τ*_*e*_ for the effective telegraph model. They are estimated to be *τ*_*c*_ = 45 min and *τ*_*e*_ = 105 min. Hence the response time gap is estimated to be *G* = *d*(*τ*_*c*_ − *τ*_*e*_) = 0.014 × (105 − 45) = 0.84, which is much larger than the maximum value of 0.67 for the negative feedback model (Fig. 4(b)). This again shows that the negative feedback model should be excluded and confirms our previous choice to use the cross-talk pathway model to interpret the data.

Our results imply that the activation of the P_lac*/*ara_ promoter is likely to be realized by the competition between a weak and a strong signalling pathway. This is supported by the biological fact that activation of the P_lac*/*ara_ promoter, under the induction of arabinose and IPTG, is regulated by both the repressor LacI and the activator AraC, which compete to bind to the promoter [1]. The unbinding of LacI from the promoter and the binding of AraC to the promoter correspond to two different pathways. The activation rate *λ*_2_ of the strong pathway is estimated to be over 40-fold larger than the activation rate *λ*_1_ of the weak pathway. The selection probabilities of the weak and strong pathways are estimated to be *q*_1_ = 0.13 and *q*_2_ = 0.87, respectively.

### Inference of gene regulation mechanisms using parameter-varying data

Experimentally, gene expression data are often measured under different experimental conditions. One of the most common experimental strategies is to modulate the value of only one parameter and keep the values of other parameters invariant, e.g. to modulate the feedback strength in a genetic feedback loop while keep the protein synthesis and decay rates, as well as the gene switching rates the same [82]. Given gene expression data under different experimental conditions, a natural question is whether we can infer the underlying gene regulation mechanisms, e.g. whether there is a feedback loop, multiple gene states, or multiple gene activation pathways.

To answer this, for each complex model, we generate synthetic data of mRNA or protein numbers using the SSA under 40 different experimental conditions, where we tune the value of only one parameter and fix the values of other parameters. The parameter that is modulated will be called the *tuning parameter* in what follows. Here we fix *λ*_1_ = 0.2 for the cross-talk pathway model so that each complex model has four tuning parameters. We choose 10 different values for each tuning parameter and hence there are 4 × 10 = 40 experimental conditions for each complex model. For each experimental condition, we then fit the steady-state simulated distribution to the telegraph model and obtain estimates of the effective parameters 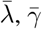, and 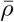.

Note that tuning a single parameter will lead to a change in the mean expression level and will also give rise to changes in the effective parameters. For each complex model and each tuning parameter, we illustrate 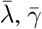, and 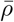 as functions of the mean expression level (Fig. 5(a)). For convenience, the value of each effective parameter is normalized to unity at the lowest mean expression level. Interestingly, when modulating a single parameter, the changes in the effective parameters follow some consistent principles. The rules for the three-state model are simple: variations in the gene activation rate *λ*_1_ (or *λ*_2_), gene inactivation rate *γ*, and synthesis rate *ρ* result in changes in their counterparts 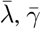, and 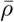 in the effective telegraph model, respectively (Fig. 5(a)). For the cross-talk pathway model, variations in *γ* and *ρ* result in changes in their counterparts 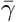 and 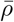, respectively, while tuning either the selection probability *q*_2_ or the gene activation rate *λ*_2_ of the strong pathway gives rise to simultaneous variations in both 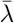 and 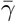 (Fig. 5(a)). Furthermore, we can distinguish between the regulations of *q*_2_ and *λ*_2_ since the increase in *q*_2_ leads to increasing 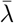 and decreasing 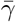, while the increase in *λ*_2_ leads to non-monotonic 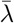 and decreasing 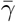.

**Figure 5:**
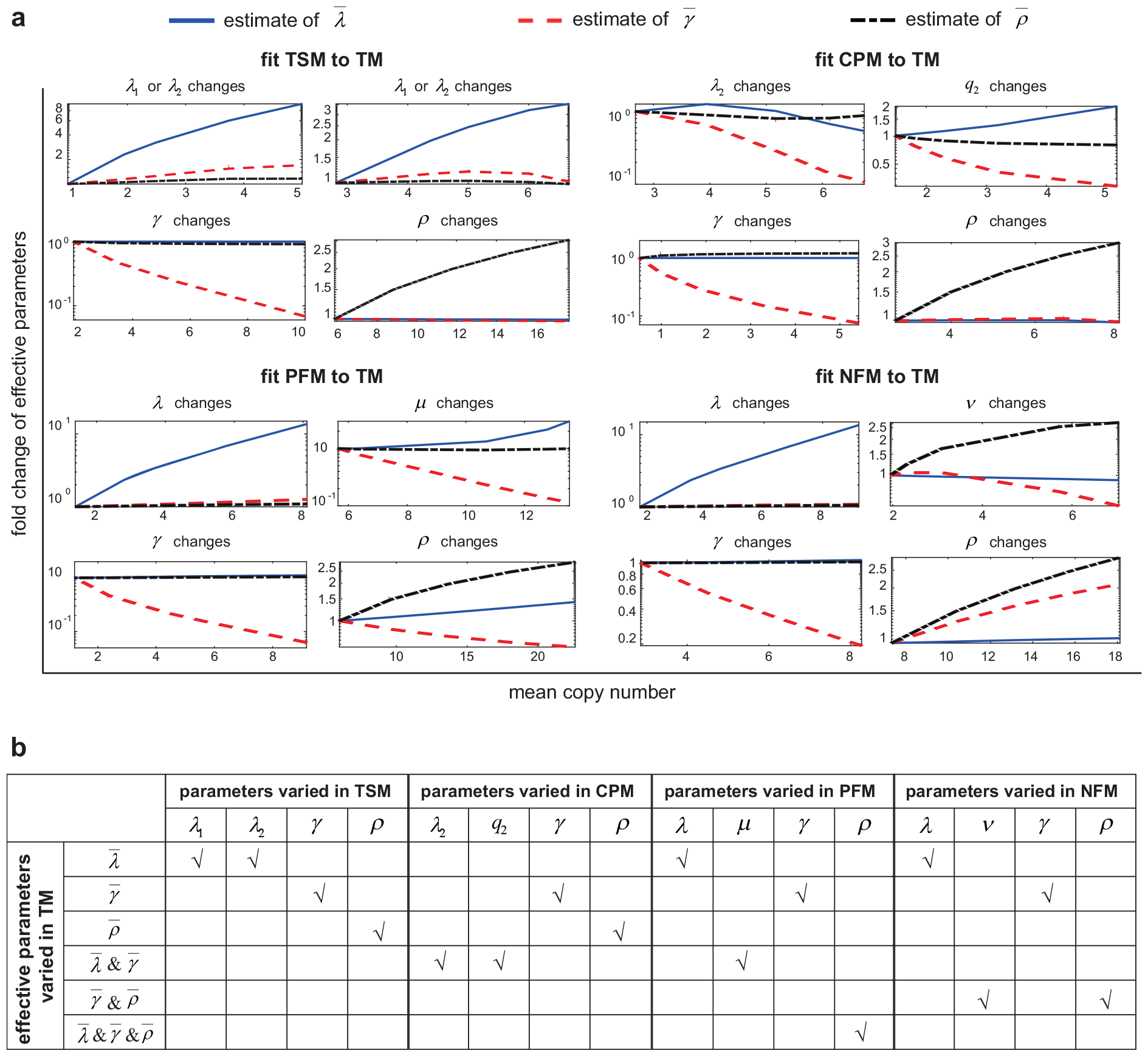
Variation patterns of effective parameters under different induction conditions in all complex models. **(a)** Tuning a single parameter of a complex model can generate a series of steady-state gene product distributions, along with different mean expression levels. Fitting these distributions to the telegraph model leads to a series of effective parameters 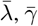, and 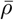. Plotting 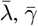, and 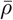 as functions of the corresponding mean expression level reveals how the effective parameters vary when a single parameter of a complex model is tuned. **(b)** Effective parameters changed when modulating a single parameter of a complex model. For example, for the positive feedback model, the effective parameter 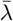 changes when tuning the parameter *λ*, while all effective parameters 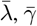, and 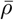 change when tuning the parameter *ρ*.

For the positive and negative feedback models, variations in the gene switching rates *λ* and *γ* result in changes in their counterparts 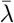 and 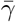 in the effective telegraph model, respectively (Fig. 5(a)). The situation is different when modulating the feedback strengths *μ* and *ν*, as well as the synthesis rate *ρ*. It is clear that the increase in the positive feedback strength *μ* leads to increasing 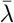 and decreasing 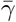; the decrease in the negative feedback strength *ν* gives rise to increasing 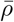 and decreasing 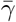; the increase in the synthetic rate *ρ* results in simultaneous increase in both 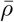 and 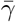 for the negative feedback model and simultaneous variations in all effective parameters 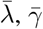, and 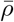 for the positive feedback model.

We emphasize that the above rules are actually independent of the choice of model parameters. To see this, for each complex model and each tuning parameter, we repeat the above procedures under 5^3^ = 125 parameter sets, where we choose five different values for each of the three fixed parameters (see Methods). All 125 parameter sets give rise to the same principles as in Fig. 5(a). For clarity, we summarize them in Fig. 5(b). These principles not only reveal the influence of the tuning parameter on the effective parameters, but also provide a potentially useful way of inferring the underlying gene regulation mechanism by using gene expression data under different induction conditions. For example, if we observe increasing *λ* (*ρ*) and decreasing *γ* as a function of the mean expression level in a gene network in response to varying experimental conditions, then we have good reasons to conjecture that there is a positive (negative) feedback loop within the network.

### Validation of theoretical results using synthetic gene networks

To validate our theory, we apply it to a synthetic gene network (orthogonal property of a synthetic network can minimize extrinsic noise) stably integrated in human kidney cells, as illustrated in Fig. 6(a) [82]. In this network, a bidirectional promoter is designed to control the expression of two fluorescent proteins: zsGreen and dsRed. The activity of the promoter can be activated in the presence of Doxycycline (Dox). The green fluorescent protein, zsGreen, is fused upstream from the transcriptional repressor LacI. The LacI protein binds to its own gene and inhibits its own transcription, forming a negative autoregulatory feedback loop. The strength of negative feedback can be tuned by induction of IPTG. As a control architecture, the red fluorescent protein, dsRed, is not regulated by induction of IPTG, forming a network with no feedback. The steady-state fluorescence intensities of zsGreen and dsRed are measured under ten different IPTG concentrations from 0 - 50 *μ*M and two different Dox concentrations (low and high) using flow cytometry.

**Figure 6:**
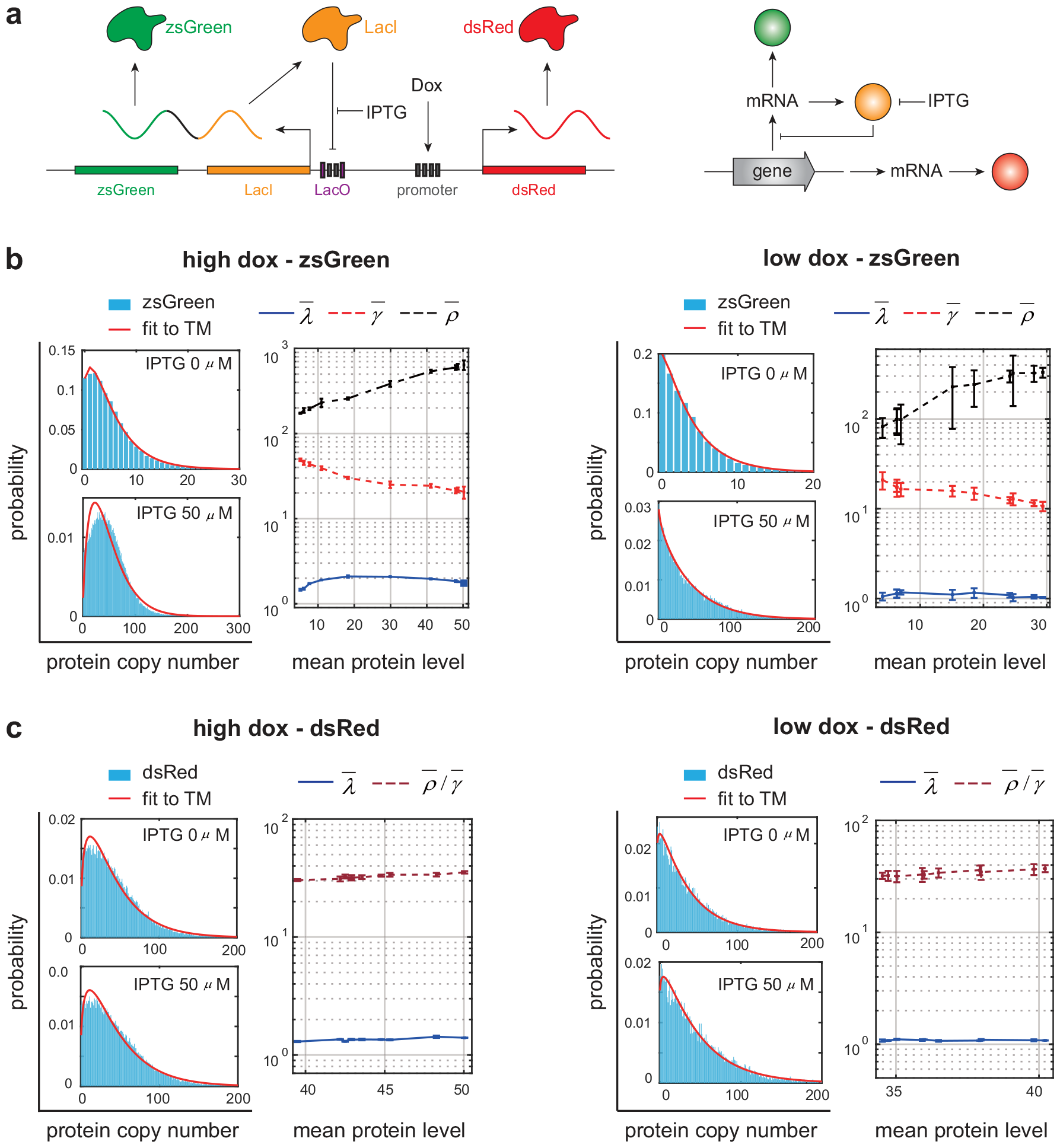
Unravelling the regulation mechanism in a synthetic gene network integrated in human kidney cells [82]. **(a)** In the network, a bidirectional promoter transcribes the zsGreen-LacI and dsRed transcripts. The gene network includes two architectures: a negative-feedback network and a network with no feedback. The zsGreen-LacI transcripts are inhibited by LacI, forming a network with negative autoregulation. The dsRed transcripts are not regulated, forming a network with no feedback. The activity of the promoter can be activated in the presence of Dox, and the negative feedback strength can be tuned by induction of IPTG. **(b)** Under both high and low Dox levels, fitting the distributions of zsGreen levels under different IPTG concentrations to the telegraph model leads to increasing 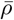, decreasing 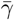, and almost invariant 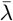 against the mean expression level. Such variation pattern of the three effective parameters coincides with that in the negative feedback model when the feedback strength *ν* is tuned. **(c)** Under both high and low Dox levels, fitting the distributions of dsRed levels under different IPTG concentrations to the telegraph model leads to almost invariant values of 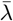 and 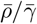 against the mean expression level. The error bars in (b) and (c) show the standard deviation of three repeated experiments [82].

Note that in this experiment, it is the fluorescence intensities of the two proteins that are measured, rather than their copy numbers. Hence it is crucial to determine the proportionality constant between fluorescence intensities and copy numbers. In other words, we need to convert the fluorescence intensity *x* into the copy number *n* = [*x/β*], where *β* represents the fluorescence intensity per protein copy and [*a*] denotes the integer part of *a*. For zsGreen, since negative feedback is weak when IPTG concentration is high, the value of *β* is chosen such that the mean number of zsGreen is equal to 50 at the highest IPTG concentration (50 *μ*M) and at high Dox concentration, which is compatible with the typical number of LacI repressor in the *lac* operon [83]. For dsRed, since its expression is not regulated by IPTG induction, the value of *β* is chosen such that the mean number of dsRed is equal to 50 at high Dox concentration.

We then fit the distributions of zsGreen levels to the telegraph model at all IPTG and Dox concentrations and obtain estimates of the effective parameters 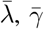, and 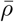. Note that increasing IPTG concentration will lead to the increase in the mean expression level. Fig. 6(b) illustrates 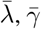, and 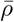 as functions of the mean expression level as IPTG concentration varies. Clearly, at both low and high Dox concentrations, the increase in IPTG concentration results in increasing 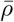 and decreasing 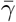, while the value of 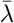 is almost unaffected by IPTG induction. This is in perfect agreement with the consistent principle shown in Fig. 5(a),(b) for the negative feedback model as the negative feedback strength *ν* changes. Hence even if we do not know in advance the topology of the network, we have good reasons to conjecture that it includes a negative feedback loop and increasing IPTG concentration weakens negative feedback. In other words, our method correctly predicts the sign of the autoregulatory loop as well as the parameter influenced by the induction conditions.

Similarly, we repeat the above procedures for dsRed. Interestingly, we find that fitting the distributions of dsRed levels to the telegraph model will lead to extremely large values of 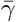 and 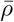, suggesting the copy number of dsRed has a negative binomial distribution (the steady-state distribution of the telegraph model reduces to a negative binomial when gene expression is sufficiently bursty, i.e. 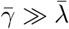and 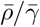 is finite [21, 84]). In this case, only the burst frequency 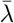 and burst size 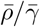 can be accurately inferred [18]. Since dsRed expression is unregulated by IPTG induction, at both high and low Dox concentrations, the values of 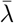 and 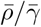 are almost invariant as IPTG concentration varies when plotted against the mean expression level (Fig. 6(c)).

### Robustness of results with respect to cooperative regulation and extrinsic noise

Note that for the feedback models shown Fig. 1(e),(f), feedback is mediated by binding of only one protein copy to the gene. However, in living systems, cooperative transcriptional regulation is very common [85]. To investigate the influence of cooperative regulation, we consider the positive and negative feedback models illustrated in Fig. 7(a), where feedback is mediated by cooperative binding of two protein copies to the gene. Again, we fit the simulated distributions obtained from the SSA to the telegraph model under 625 parameter sets and obtain estimates of the effective parameters 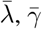, and 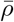.

**Figure 7:**
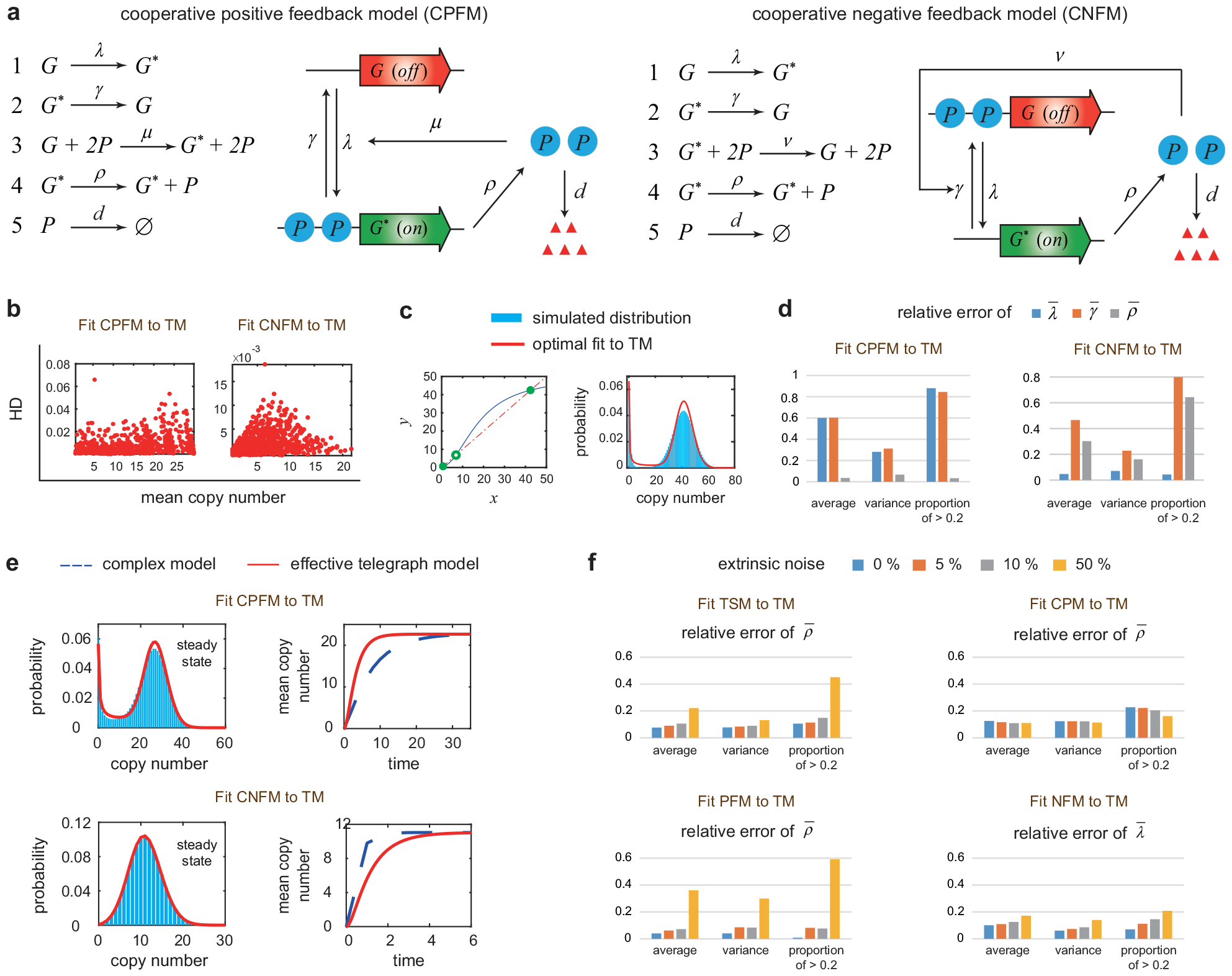
Robustness of results with respect to cooperative regulation and extrinsic noise. **(a)** Positive and negative feedback models with cooperative regulation. Feedback is mediated by cooperative binding of two protein copies to the gene. **(b)** For each cooperative feedback model, the HD between the simulated distribution and its telegraph model approximation is shown as a function of the mean expression level for 625 parameter sets. The simulated distribution is well captured by the telegraph model, manifested by HD *<* 0.065. **(c)** Under cooperative regulation and fast gene switching, the deterministic rate equation for the positive feedback model is given by 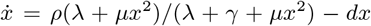. It may have two stable fixed points (and an unstable fixed point) and thus gives rise to deterministic bistability. The intersections of *y* = *ρ*(*λ* + *μx*^2^)*/*(*λ* + *γ* + *μx*^2^) (blue curve) and *y* = *dx* (red dashed curve) give the locations of the three fixed points (green circles). For a positive feedback loop with deterministic bistability, the effective telegraph model still accurately captures the resulting bimodal distribution. The parameters of the positive feedback model are chosen as *ρ* = 50, *d* = 1, *λ* = 2, *γ* = 160, *μ* = 0.5. The effective parameters are estimated to be 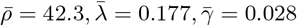. **(d)** For each cooperative feedback model, the (absolute values of) relative errors of the three effective parameters 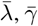 and 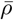 are computed under 625 parameter sets, along with their sample means, sample variances, and the sample frequencies of relative errors being greater than 0.2. **(e)** Under cooperative regulation, the positive feedback model still has smaller time-dependent mean curve compared to the effective telegraph model, while the negative feedback model still has larger time-dependent mean curve. **(f)** For each complex model and each parameter set, the simulated distributions are fitted to the telegraph model under four noise levels (0%, 5%, 10%, and 50%). The relative errors of the three effective parameters are computed for all parameter sets, along with the three statistics of relative errors (same as in (d)). The three statistics of 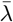 are shown for the negative feedback model, and the three statistics of 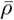 are shown for the other three complex models.

We find that almost all results obtained previously remain unchanged. First, under cooperative regulation, the steady-state protein distributions for the feedback models are still well fitted by the effective telegraph model, manifested by low HD values (Fig. 7(b)). The only difference is that in the presence of cooperative binding, the positive feedback model may produce deterministic bistability, which means that the deterministic rate equation for the system may have two stable fixed points (Fig. 7(c), left panel) [86]; this can even happen when the gene switches very rapidly between the two states, i.e. *λ* + *μ ⟨n⟩* ^2^, *γ ≫ ρ, d*. Interestingly, for a positive feedback loop with deterministic bistability, the effective telegraph model still accurately reproduces the resulting bimodal distribution by setting very small effective gene switching rates 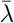and 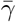 (Fig. 7(c), right panel). This coincides with our previous finding that 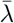 and 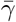 are both under-estimated in the positive feedback model.

Second, under cooperative regulation, fitting the steady-state distribution to the telegraph model yields reliable estimation of the synthesis rate *ρ* for the positive feedback model and reliable estimation of the gene activation rate *λ* for the positive feedback model (Fig. 7(d)). Comparing Fig. 3(a) with Fig. 7(d), we find that the inference of *λ* is even more accurate in the presence of cooperative regulation — the mean relative error of 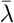 is 0.13 for the non-cooperative case and is only 0.04 for the cooperative case. Third, the time-dependent mean curve for the positive (negative) feedback model is still below (above) its counterpart for the effective telegraph model due to slower (faster) relaxation speed to the steady state (Fig. 7(e)). Finally, the variation patterns of the three effective parameters under different induction conditions also remain unchanged (Supplementary Fig. S6).

Thus far, we only consider models with intrinsic noise (Fig. 1(a)). However, extrinsic noise may contribute substantially to the gene product fluctuations, especially when intrinsic noise is small [87]. Extrinsic noise may have various sources such as transcription factor concentrations, RNA polymerase number, cellular volume, and local cell crowding [69]. A recent study [88] found that in the presence of extrinsic noise, fitting gene expression data to the standard telegraph model may lead to inaccurate parameter inference. We next investigate how the conclusions of the present paper are affected by extrinsic noise. To characterize extrinsic noise, following [77, 88], we add noise to the synthesis rate *ρ* for all complex models. Specifically, we reset *ρ* in each complex model as a log-normal distributed random variable with its mean being the original value of *ρ* and standard deviation being equal to 0.05, 0.1, and 0.5 of the mean, corresponding to noise levels of 5%, 10%, and 50%, respectively.

For each complex model and each noise level, we fit the simulated distributions obtained from the SSA to the telegraph model under 625 parameter sets. We find that the almost all results obtained previously remain unchanged when the noise level is below 10%, but some results may be broken when the noise level is increased to 50%. First, the telegraph model can still accurately reproduce the steady-state gene product distribution for all noise levels, manifested by low HD values, although the HD increases slightly with respect to the noise level (Supplementary Fig. S7). Second, for a noise level less than 10%, the gene activation rate *λ* can still be accurately estimated for the negative feedback model and the synthesis rate *ρ* can still be accurately estimated for the other three complex models, similarly to models without extrinsic noise (Fig. 7(f)). When the noise level is increased to 50%, the estimate of *ρ* is still accurate for the cross-talk pathway model and the estimate of *λ* is still accurate for the negative feedback model; however, there is a sharp increase in the mean relative error for the other two complex models. Interestingly, combining Fig. 7(e),(f), we find that the robustness of parameter inference with respect to extrinsic noise for a given model is closely related to its relaxation speed to the steady state — faster relaxation speed results in more precise inference under large extrinsic noise.

Third, for all noise levels, the three-state and positive feedback models still have slower relaxation speed to the steady state compared to the effective telegraph model, while the cross-talk pathway and negative feedback models relax to the steady state faster (Supplementary Fig. S8). Finally, the variation patterns of the three effective parameters under different induction conditions remain unchanged for small and intermediate noise levels and may change dramatically when the noise level is increased to 50% (Supplementary Fig. S9-S12). In summary, all the conclusions in the present paper are robust in the presence of cooperative regulation and (small or intermediate) extrinsic noise.

## Conclusions and discussion

A central question in molecular biology is to understand various genetic regulation mechanisms and how they modulate the production of mRNA and protein at the single-cell level [28, 47]. The classical telegraph model has been extensively used to explain single-cell gene expression data so that one can estimate the underlying kinetic parameters and unravel gene regulation mechanisms in response to varying environmental changes [15, 17, 29]. However, the telegraph model is limited in its predictive power since it lacks a description of some biological mechanisms that are known to have a profound impact on the mRNA and protein distributions in single cells. In the presence of complex biological mechanisms, fitting gene expression data to the simple telegraph model [17] may lead to inaccurate parameter inference and even incorrect predictions of the underlying gene regulation mechanisms.

In the present paper, we investigate the dynamical properties of four relatively complex gene expression models, including the three-state, cross-talk pathway, positive feedback, and negative feedback models. Compared with the telegraph model, these models describe how fluctuations are influence by complex biological mechanisms such as non-exponential gene inactivation durations, multiple gene activation pathways, and feedback regulation. Our method is to fit the steady-state mRNA or protein distribution of each complex model to a simple telegraph model for a large sets of model parameters. Despite the potential risks, we found that fitting these complex models to the telegraph model still provide a large amount of valuable information. In fact, the idea of using the distribution of the telegraph model to approximate that of a complex model has been applied in previous studies using analytical methods such as linear mapping or moment matching [89–91]. Here we evaluate the performance of the effective telegraph model using statistical and computational methods.

First, we showed that the steady-state gene product distributions, as well as the conditional distributions in the active gene state, of the four complex models can all be well fitted by the telegraph model. We found that while most effective parameters may deviate significantly from their real values in the complex models, there are still some parameters that can be reliably estimated with very small relative errors. For the three-state, cross-talk pathway, and positive feedback models, the effective synthesis rate is very closed to its real value, while the effective gene activation and inactivation rates deviate largely from their real values. At first glance, only the gene activation mechanism in the three complex models differs from that in the telegraph model. However, our results showed that fitting the steady-state distributions of the complex models to the telegraph model may lead to unreliable estimation of both the gene activation and inactivation rates [7], but does not significantly influence the synthesis rate. For the negative feedback model, we showed that the effective synthesis and gene inactivation rates are unreliable, while the effective gene activation rate is very closed to its real value.

The effective parameters also provide a natural and convenient way of characterizing the capability for a complex model to exhibit bimodal gene product distributions. This characterization is based on a mathematical result [35, 92] which shows that the telegraph model can generate a bimodal distribution only when its gene activation and inactivation rates are both smaller than the decay rate. For the three-state model, the effective gene switching rates are both over-estimated compared to their real values, which makes bimodality difficult to occur. In contrast, for the cross-talk pathway and positive feedback models, the effective gene switching rates are both under-estimated, and thus these two models are more likely to exhibit bimodality. For the negative feedback model, one of the effective gene switching rates is over-estimated while the other is under-estimated, which exerts a weak influence on bimodality.

Furthermore, we showed that the effective parameters can be used to distinguish a complex model from the telegraph model by using additional single-cell data at multiple time points. The good fit of complex models to the telegraph model in steady state indicates that it is impossible to distinguish between the two models by only using the steady-state gene expression data. Previous studies showed that a non-monotonic dynamic feature of gene expression mean can rule out the telegraph model since the telegraph model can only display a monotonic time-dependent mean curve [56, 57]. However, this does not work when the gene expression mean displays a monotonic dynamics. To solve this, we compared the time-dependent mean curves of a complex model and its effective telegraph model, where the effective parameters were estimated using the steady-state data. We showed that if the mean curve for the effective telegraph model is below the real mean curve, then the three-state or positive feedback model is more competitive to describe the data compared to the telegraph model. In contrast, if the former is above the latter, we may select the cross-talk pathway or negative feedback model to explain the data. A method based on the response times of a complex model and its effective telegraph model can further distinguish the cross-talk pathway model from the negative feedback model. As a validation of our method, we apply it to the snapshot mRNA expression data of the P_lac*/*ara_ promoter at multiple time points measured in *E. coli* cells [1]. We showed that among the four complex models, the cross-talk pathway model is the most competitive and thus we predict that the activation of P_lac*/*ara_ is very likely to be regulated by the competition between two signalling pathways.

In addition, we showed that the effective parameters can be used to unravel the gene regulation mechanism of a complex model in response to varying environmental conditions. For a series of gene product distributions obtained by tuning a single parameter of a complex model, fitting those distributions to the telegraph model gives a certain variation pattern of the three effective parameters. We found that the variation pattern of effective parameters is independent of the choice of parameters of the complex model. Hence it is possible to determine the underlying gene regulatory mechanism in response to environmental changes by identifying the variation pattern of effective parameters. To test our method, we apply it to the protein expression data of a synthetic autoregulatory gene circuit in human kidney cells which is designed to suppress gene expression under IPTG induction [82]. Fitting the steady-state data under all induction conditions to the telegraph model reveals a certain variation pattern of effective parameters, which is in perfect agreement with that of the negative feedback model by tuning the negative feedback strength. Hence our method correctly predicts the sign of the autoregulatory loop as well as the parameter influenced by the induction conditions. In contrast, fitting the data for an unregulated system to the telegraph model results in almost constant effective parameters as IPTG concentration varies.

Finally, we showed that almost all results in the present paper are robust with respect to cooperative transcriptional regulation and extrinsic noise. In particular, we find that the robustness of parameter inference with respect to extrinsic noise for a given model is closely related to its relaxation speed to the steady state — faster relaxation speed results in more precise inference under large extrinsic noise.

In summary, the telegraph model should be used with caution when there are complex mechanisms behind the underlying gene expression system. However, fitting the steady-state distribution of a relatively complex gene expression model to the telegraph model can still reveal rich information. Specifically, we learn that (i) some effective parameters are reliable and can reflect realistic dynamic behavior of the complex model; (ii) the under-estimation of the effective gene activation and inactivation rates reveals the ability for a complex model to exhibit bimodality; (iii) time-resolved data are needed to distinguish between different mechanisms; (iv) comparing the time evolution of the mean expression level with the prediction of the effective telegraph model provides a method of reliable model selection; (v) the variation pattern of effective parameters can reveal gene regulation mechanisms in response to environmental changes.

The current study has some limitations. First, in our feedback models, we assume that there is no change in the protein number during gene activation and inactivation. However, in reality, the protein number decreases by one when a protein copy binds to a gene and increases by one when unbinding occurs [43]. Here we make this assumption because it leads to a simple analytical expression of the protein distribution so that the mean active and inactive durations can be solved exactly [40, 93]. Second, in our feedback models, we ignore the mRNA dynamics and assume that protein is produced directly from the gene. This is a reasonable simplification when mRNA decays much faster than protein and the burst size of protein is relatively small [21, 93]. Here we make this assumption because incorporating the mRNA description into the feedback models leads to two additional parameters which will complicate theoretical analysis and parameter inference. Last but not least, while our model takes extrinsic noise into account, it does not incorporate post-transcriptional sources of noise such as RNA splicing and nuclear export, which affect mature mRNA but not nascent mRNA. Recent studies [69] have shown that parameters estimated using the telegraph model or other models relying on mature mRNA may be suspicious because they differ from those estimated using nascent mRNA data. This cannot be removed even with time-dependent modelling unless one explicitly models post-transcriptional noise.

Future work is required to further test our methods by adding more detailed biological mechanisms into the telegraph model, including bursty production of mRNA and protein [94, 95], cell cycle events such as cell growth and division [96, 97], cell-volume dependence [22, 98], as well as complex gene regulatory networks [32, 89]. In addition, we anticipate that our theory can be enriched by fitting the time-dependent distributions (rather than only the steady-state distributions) of complex models to the telegraph model. For complex models, a detailed comparison between the maximum-likelihood method and other parameter inference methods such as Bayesian inference is also expected.

## Methods

### Selection of parameter sets for complex models

In Fig. 2, we generate synthetic data of gene product numbers under 625 different parameter sets for the four complex models. For convenience, we set *d* = 1 for all models. For the three-state model, we set *ρ* = 10, 15, 20, 25, 30 and *λ*_1_, *λ*_2_, *γ* = 0.3, 0.7, 1, 2, 4, which gives 5^4^ = 625 combinations of the four parameters. For the cross-talk pathway model, the gene activation rate for the weak signalling pathway is fixed to be *λ*_1_ = 0.2. The other four parameters are chosen as *ρ* = 10, 15, 20, 25, 30, *λ*_2_, *γ* = 0.5, 1, 2, 4, 8, and *q*_1_ = 0.1, 0.3, 0.5, 0.7, 0.9. For the positive and negative feedback models, we set *ρ* = 10, 15, 20, 25, 30, *λ, γ* = 0.3, 0.7, 1, 2, 4, and *μ, ν* = 0.05, 0.1, 0.5, 1, 1.5.

In Fig. 3(c), we randomly select 150 parameter sets such that the values of 1*/⟨T*_off_*⟩* and 1*/⟨T*_on_*⟩* are between 0 and 2.5*d* for each complex model. The synthesis rate *ρ* is randomly selected so that *ρ* ∈ [10, 30]. For the three-state, cross-talk pathway, and positive feedback models, 1*/⟨T*_on_*⟩* = *γ* is randomly selected so that *γ* ∈ [0.1, 2.5]. For the negative feedback model, 1*/⟨T*_off_*⟩* = *λ* is randomly selected so that *λ* ∈ [0.1, 2.5]. Moreover, for the three-state model, the gene activation rates *λ*_1_ and *λ*_2_ are randomly selected so that *λ*_1_, *λ*_2_ ∈ [0.1, 5]. This ensures that

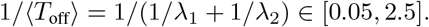

For the cross-talk pathway model, the selection probability *q*_1_ is randomly selected so that *q*_1_ ∈ [0.1, 0.9] and the gene activation rates *λ*_1_ and *λ*_2_ for the two pathways are randomly selected so that*λ*_1_ ∈ [0.05, 0.5] and *λ*_2_ ∈ [1, 8]. This ensures that

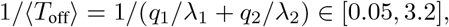

and we randomly select 150 parameter sets that satisfy 1*/⟨T*_off_*⟩* ≤ 2.5. The analytical formula of *⟨T*_off_*⟩* for the positive feedback model and the analytical formula of *⟨T*_on_*⟩* for the negative feedback model are too complicated to directly calculate their upper and lower bounds. To overcome this, we restrict *λ* ∈ [0.1, 1.5] and *μ* ∈ [0.01, 0.15] for the positive feedback model and randomly select 150 parameter sets that satisfy 1*/⟨T*_off_*⟩* ≤ 2.5. Similarly, we restrict *γ* ∈ [0.1, 1.5] and *ν* ∈ [0.01, 0.15] for the negative feedback model and randomly select 150 parameter sets that satisfy 1*/⟨T*_on_*⟩* ≤ 2.5.

In Fig. 5, we tune a single parameter while fix the other parameters for each complex model. The values for parameters are chosen to be the same as in Fig. 2. Hence each complex model has four parameters to be tuned and each tuning parameter is equipped with five different values. For each complex model, we tune a single parameter among 10 different values, and hence there are total 5^3^ = 125 combinations for the other three parameters. To observe a significant effect of the tuning parameter on the mean gene expression level, we only consider those combinations of the other three parameters such that the mean expression level changes by at least two folds.

### Computation of the confidence interval

For the cross-talk pathway model, we use the profile likelihood method [17] to compute the confidences intervals of all parameters. For example, for the parameter *λ*_1_, we start by fixing *λ*_1_ and vary all other parameters to maximize the log-profile-likelihood function

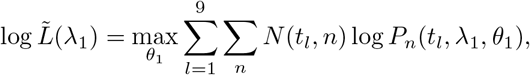

where *θ*_1_ is the freely varying parameter set (*λ*_2_, *q*_1_, *q*_2_, *γ, ρ*) for the cross-talk pathway model and the meanings of other quantities are the same as in Eq. (3). When the sample size is large, the statistics

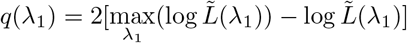

asymptotically approaches the chi-square distribution 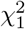 with one degree of freedom [17]. The point estimate of *λ*_1_ must satisfy *q*(*λ*_1_) = 0. To obtain the 95% confidence interval, we find the parameter region of *λ*_1_ such that

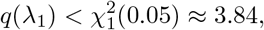

where 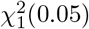 is the cutoff value for 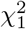 under the significance level of 0.05.

## Author contributions

C. J. conceived the original idea. F. J. and C. J. performed the theoretical derivations. F. J., J. L., T. L., Y. Z., and W. C. perform the numerical simulations and data analysis. F. J., L. B., and C. J. interpreted the theoretical results. F. J. and C. J. jointly wrote the manuscript.

## Data and materials availability

All data needed to evaluate the conclusions are present in the paper. The MATLAB codes for parameter inference can be found on GitHub via the link http://cam.gzhu.edu.cn/info/1014/1223.htm or the link https://github.com/chenjiacsrc/telegraph-model-fitting.

## Acknowledgements

F. J. acknowledges support from National Natural Science Foundation of China with grant No. 12271118. C. J. acknowledges support from National Natural Science Foundation of China with NSAF grant No. U2230402 and grant No. 12271020.

