## Supplementary for "What can we learn when fitting a simple telegraph model to a complex gene expression model?"

#### Contents

**1 Analytical distributions for the four complex models** **2**

**2 Parameter inference for the telegraph model** **2**

**3 Mean active and inactive periods for feedback models** **3**

### 1 Analytical distributions for the four complex models

Let $P_n$ denote the steady-state probability of having $n$ copies of the gene product in a single cell. For the three-state model, the corresponding generating function $F(z) = \sum_{n=0}^{\infty} P_n z^n$ of the gene product distribution is given by [1]

$$F(z) = {}_2F_2(\lambda_1, \lambda_2; \alpha_1, \beta_1; \rho(z-1)), \tag{1}$$

where $\alpha_1 + \beta_1 = \lambda_1 + \lambda_2 + \gamma$, $\alpha_1\beta_1 = \lambda_1\lambda_2 + \lambda_1\gamma + \lambda_2\gamma$, and ${}_2F_2$ denotes the generalized hypergeometric function. Once the generating function $F(z)$ is known, the gene product distribution $P_n$ can be recovered by taking the derivative of $F(z)$ at $z = 0$:

$$P_n = \frac{1}{n!}\frac{d^nF(z)}{dz^n}\bigg|_{z=0}.$$

For the cross-talk pathway model, the generating function of the gene product distribution is given by [2]

$$F(z) = {}_2F_2(\lambda_1, \lambda_2; \alpha_2, \beta_2; \rho(z-1)), \tag{2}$$

where $\alpha_2 + \beta_2 = \lambda_1 + \lambda_2 + \gamma$ and $\alpha_2\beta_2 = \lambda_1\lambda_2 + (q_2\lambda_1 + q_1\lambda_2)\gamma$.

For the positive feedback model, the The generating function of the gene product distribution is given by [3]

$$F(z) = \frac{{}_1F_1(\alpha_3; \beta_3; \rho(z - z_3))}{{}_1F_1(\alpha_3; \beta_3; \rho(1 - z_3))}, \tag{3}$$

where $\alpha_3 = \lambda/(\mu + 1)$, $\beta_3 = (\lambda + \gamma)/(\mu + 1)$, $z_3 = 1/(\mu + 1)$, and ${}_1F_1$ denotes the confluent hypergeometric function.

For the negative feedback model, the generating function of the gene product distribution is given by [4]

$$F(z) = \frac{{}_1F_1(\alpha_4; \beta_4; \rho(z - z_4))}{{}_1F_1(\alpha_4; \beta_4; \rho(1 - z_4))}, \tag{4}$$

where $\alpha_4 = (\lambda + \gamma)/(\nu + 1) + \rho\nu/(\nu + 1)^2$, $\beta_4 = \rho/(\nu + 1)$, and $z_4 = 1/(\nu + 1)$.

### 2 Parameter inference for the telegraph model

We use the "fminsearch" function in MATLAB to minimize the log-likelihood function

$$\log L(\theta) = \sum_n N(n)\log(P_n(\theta)), \tag{5}$$

where $\theta$ is the parameter set, $N(n)$ denotes the number of cells with $n$ copies of mRNA or protein, and $P_n(\theta)$ denotes the mRNA or protein number distribution with parameter set $\theta$. In this way, we can obtain estimates of the three parameters $\bar{\lambda}$, $\bar{\gamma}$, and $\bar{\rho}$ of the telegraph model (assuming that $d = 1$). One of the most important steps for optimization is to select the initial values $\bar{\lambda}_0$, $\bar{\gamma}_0$, and $\bar{\rho}_0$ of the three parameters. The conventional moment estimation method determines $\bar{\lambda}_0$, $\bar{\gamma}_0$, and $\bar{\rho}_0$ as functions of the first three moments of gene expression data across a cell population by solving three algebraic equations [5, 6]. Although this method is straightforward, it may produce negative values of $\bar{\lambda}_0$, $\bar{\gamma}_0$, and $\bar{\rho}_0$ [6]. To avoid the possible negative initial values, we modify the moment estimation method as follows. If $\bar{\rho}_0$ is known, then the two initial values $\bar{\lambda}_0$ and $\bar{\gamma}_0$ can be determined by matching the mean and variance of gene product fluctuations [7, 8], which leads to

$$\bar{\lambda}_0 = \frac{M(\bar{\rho}_0 + 1 - \eta M - M)}{\bar{\rho}_0(\eta M - 1)}, \quad \bar{\gamma}_0 = \frac{\bar{\rho}_0 - M}{M}\bar{\lambda}_0, \tag{6}$$

where $M$ is the mean copy number and $\eta$ is the coefficient of variation squared. Note that $\bar{\rho}/d$ is the typical gene product number in the active gene state. Hence in previous papers [7], $\bar{\rho}_0$ is commonly chosen as the maximum number of gene product molecules within the cell population. However, this choice may still give rise to negative values of $\bar{\lambda}_0$ and $\bar{\gamma}_0$ in Eq. (6) when the maximum number of molecules $\bar{\rho}_0 < \eta M + M - 1$. If this happens, then we simply reset $\bar{\rho}_0 = \eta M + M$. To test this method, we use the telegraph model to generate synthetic data of gene products numbers for $N = 10^5$ cells using the SSA under $10^3$ randomly selected parameter sets within the regions $\bar{\lambda}, \bar{\gamma} \in [0.1, 4]$ and $\bar{\rho} \in [10, 30]$. The synthetic data are then fitted to the telegraph model using the maximum-likelihood method with the initial values $\bar{\lambda}_0$, $\bar{\gamma}_0$, and $\bar{\rho}_0$ being chosen as above. For all $10^3$ parameter sets, the estimated values of $\bar{\lambda}$, $\bar{\gamma}$, and $\bar{\rho}$ are almost the same as their real values (Supplementary Fig. S5). This shows that our parameter inference method is highly reliable.

The readers may also ask how frequently the resetting of $\bar{\rho}_0$ occurs for real data. To see this, we generate synthetic data for $N = 10^2, 10^3, 10^4, 10^5$ cells under $10^3$ randomly selected parameter sets within the same regions mentioned above. Interestingly, we find that the resetting of $\bar{\rho}_0$ fails to be observed for all the tested sample sizes and parameter sets. This indicates that such resetting should rarely occur for real smFISH or scRNA-seq data.

### 3 Mean active and inactive periods for feedback models

We next compute the the mean active and inactive durations for the two feedback models. In [3], it has been shown that no matter there is autoregulation or not, the mean of gene product numbers can always be represented by the mean active and inactive periods as

$$\langle n \rangle = \rho \times \frac{\langle T_{\text{on}} \rangle}{\langle T_{\text{on}} \rangle + \langle T_{\text{off}} \rangle}, \tag{7}$$

where $\langle T_{\text{on}} \rangle$ and $\langle T_{\text{off}} \rangle$ are the mean active and inactive periods of the gene, and the second term on the right-hand side represents the probability of the gene being active. This formula is intuitive for unregulated genes. However, for regulated genes, it is highly non-trivial and was proved in [3] using the analytical distributions of gene product numbers. The mean protein number $\langle n \rangle = F'(1)$ can be easily recovered by taking the derivatives of the generating function given in Eqs. (3) and (4) at $z = 1$. For the positive feedback model, the gene expression mean $\langle n \rangle$ has the form of [3]

$$\langle n \rangle = \rho \times \frac{\alpha_3 \cdot {}_1F_1(\alpha_3 + 1; \beta_3 + 1; \rho(1 - z_3))}{\beta_3 \cdot {}_1F_1(\alpha_3; \beta_3; \rho(1 - z_3))}.$$

Moreover, the mean active duration for the positive feedback model is given by $\langle T_{\text{on}} \rangle = 1/\gamma$. Substituting this equation into Eq. (7) gives the explicit expression of the mean inactive duration, i.e.

$$\langle T_{\text{off}} \rangle = \frac{\beta_3 \cdot {}_1F_1(\alpha_3; \beta_3; \rho(1 - z_3)) - \alpha_3 \cdot {}_1F_1(\alpha_3 + 1; \beta_3 + 1; \rho(1 - z_3))}{\gamma \alpha_3 \cdot {}_1F_1(\alpha_3 + 1; \beta_3 + 1; \rho(1 - z_3))}.$$

Similarly, for the negative feedback model, the gene expression mean is given by [3]

$$\langle n \rangle = \rho \times \frac{\alpha_4 \cdot {}_1F_1(\alpha_4 + 1; \beta_4 + 1; \rho(1 - z_4))}{\beta_4 \cdot {}_1F_1(\alpha_4; \beta_4; \rho(1 - z_4))}.$$

Moreover, the mean inactive duration for the negative feedback model is given by $\langle T_{\text{off}} \rangle = 1/\lambda$. Substituting this equation into Eq. (7) gives the explicit expression of the mean active duration, i.e.

$$\langle T_{\text{on}} \rangle = \frac{\alpha_4 \cdot {}_1F_1(\alpha_4 + 1; \beta_4 + 1; \rho(1 - z_4))}{\lambda \beta_4 \cdot {}_1F_1(\alpha_4; \beta_4; \rho(1 - z_4)) - \lambda \alpha_4 \cdot {}_1F_1(\alpha_4 + 1; \beta_4 + 1; \rho(1 - z_4))}.$$

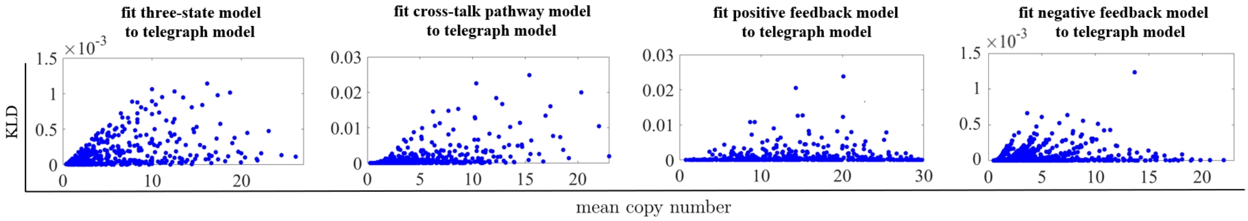


Figure 1: **Kullback-Leiber divergence (KLD) for quantifying the fit of the gene product distributions of complex models to the telegraph model.** For each complex model, synthetic data of gene product numbers are generated under 625 parameter sets using the SSA. For each complex model, the KL divergence between the simulated distribution and its telegraph model approximation is shown as a function of the mean expression level for the 625 parameter sets. The KL divergence is less than 0.025 for all complex models.

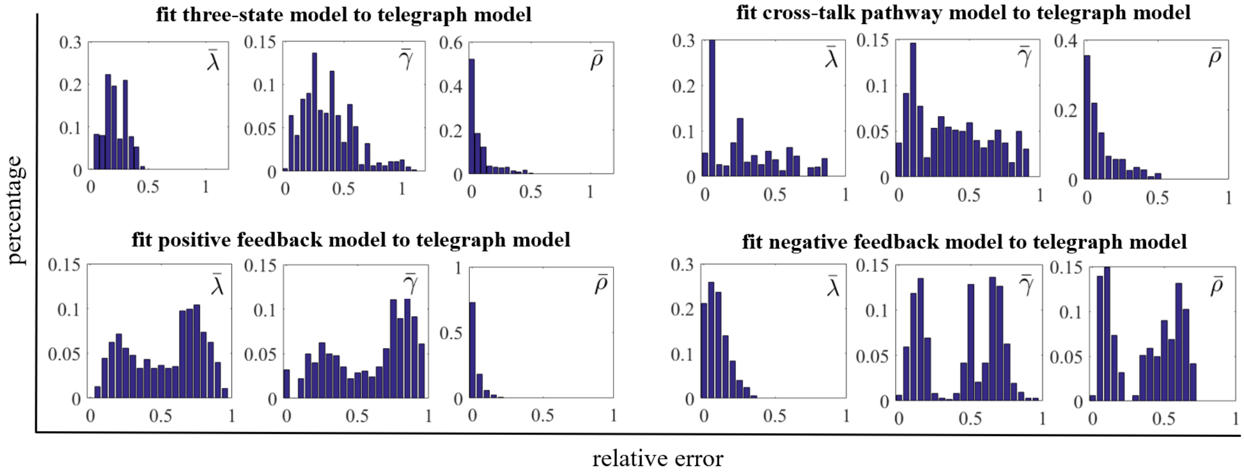


Figure 2: **Empirical distributions of the relative errors for the effective rate parameters.** For each complex model, the empirical distributions of relative errors of the three effective rates $\bar{\lambda}$, $\bar{\gamma}$ and $\bar{\rho}$ are computed under 625 parameter sets which are chosen to be the same as in Fig. 2 in the main text. For the three-state, cross-talk pathway and positive feedback models, the distribution bars for the relative errors of $\bar{\lambda}$ and $\bar{\gamma}$ scatter within a broad region, while the most relative errors of $\bar{\rho}$ are relatively small. The situation is different for the negative feedback model, for which the distribution bars for the relative errors of $\bar{\gamma}$ and $\bar{\rho}$ scatter within a broad region, and the most relative error values of $\bar{\lambda}$ are relatively small.

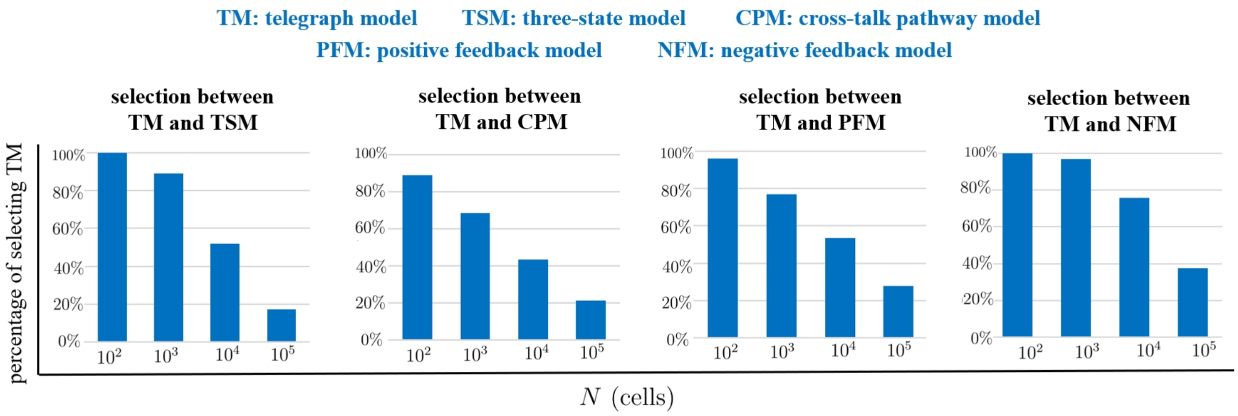


Figure 3: **Steady-state distribution data is unable to reliably distinguish a complex model from the telegraph model.** For each complex model, we generate synthetic distribution data with $N = 10^2, 10^3, 10^4, 10^5$ cells under 625 parameter sets, and then fit the synthetic data to the complex model and effective telegraph model by maximizing the likelihood. To penalize for the parameter number for complex models, we compute the corrected Akaike information criterion (AICc) and select the best model corresponding to the smallest AICc. It shows that the proportion of wrong model selection (selecting telegraph model) decreases with respect to cell sample size $N$: there are over 90% parameter sets corresponding to the wrong selection when $N = 10^2$, and over 20% parameter sets corresponding to the wrong selection for the largest $N = 10^5$.

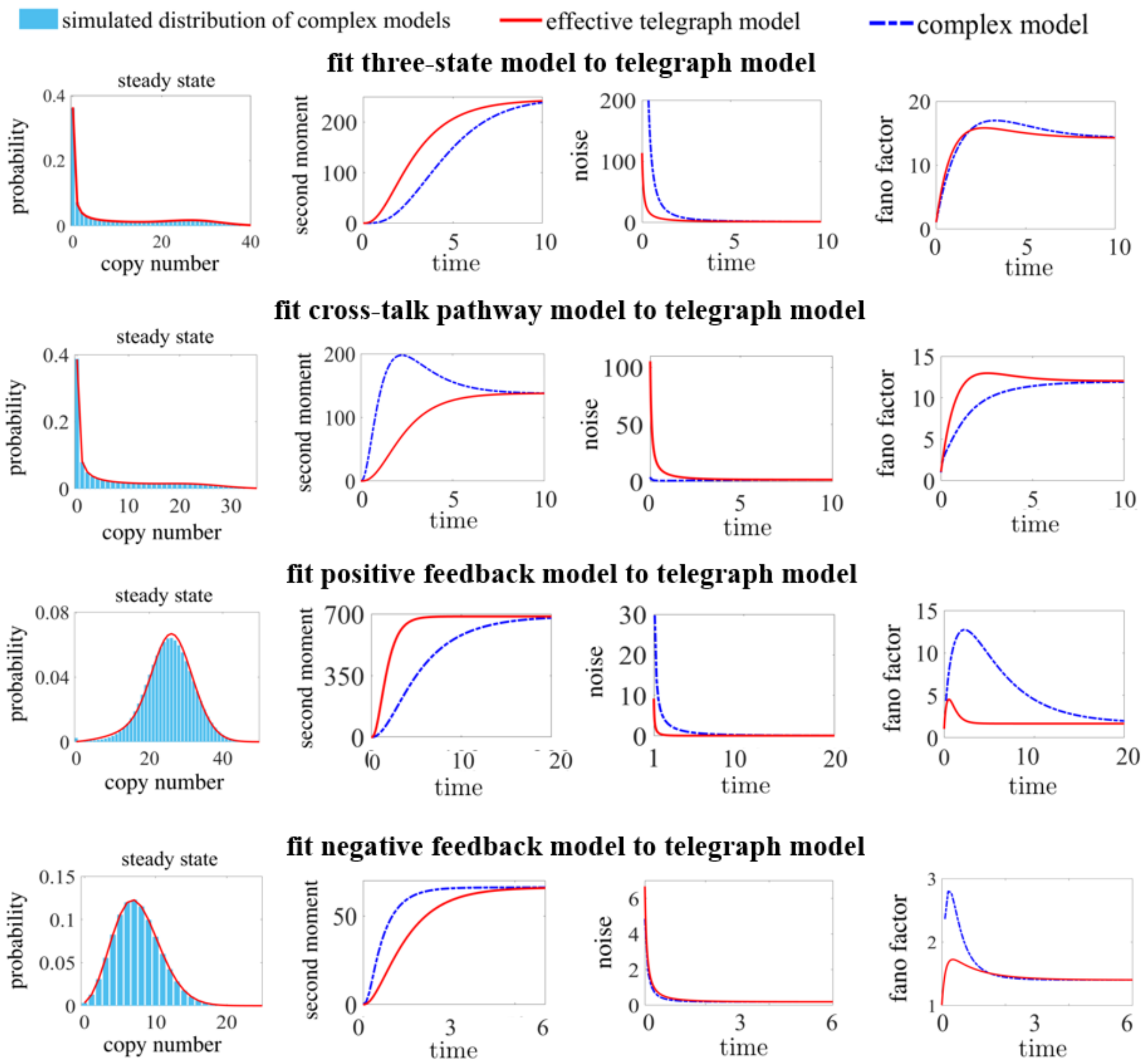


Figure 4: **Distinguishing the dynamic differences between a complex model and its effective telegraph model.** Under the initial condition that the gene is off and there is no gene product molecules in the cell, the three-state and positive feedback models have smaller time-dependent second moment curve compared to the effective telegraph model, while the cross-talk pathway and negative feedback models have larger time-dependent second moment curve. However, common fluctuation indicators such as gene expression noise and the Fano factor, present less easily distinguishable dynamic differences between complex models and their effective telegraph models.

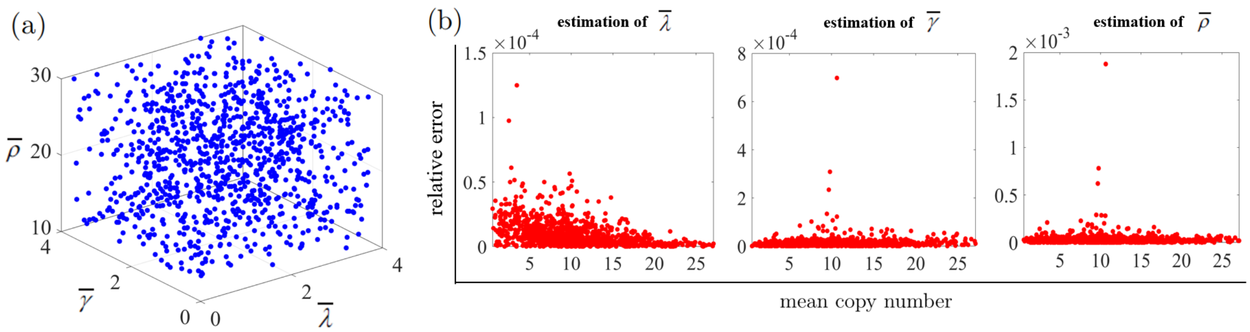


Figure 5: **High accuracy of the method for inferring parameters of the telegraph model.** **(a)** $10^3$ parameter sets of $(\bar{\lambda}, \bar{\gamma}, \bar{\rho})$ are randomly selected within the regions $\bar{\lambda}, \bar{\gamma} \in [0.1, 4]$ and $\bar{\rho} \in [10, 30]$. **(b)** For each of $10^3$ parameter sets, the synthetic distribution data is fitted to the telegraph model using the maximum-likelihood method, where the initial values $\bar{\lambda}_0$, $\bar{\gamma}_0$, and $\bar{\rho}_0$ for optimization are chosen by our method in the main text (see details in Method). The estimated values of $\bar{\lambda}$, $\bar{\gamma}$, and $\bar{\rho}$ are almost as the same as their real values, quantified by very small relative errors between real and inferred parameter values.

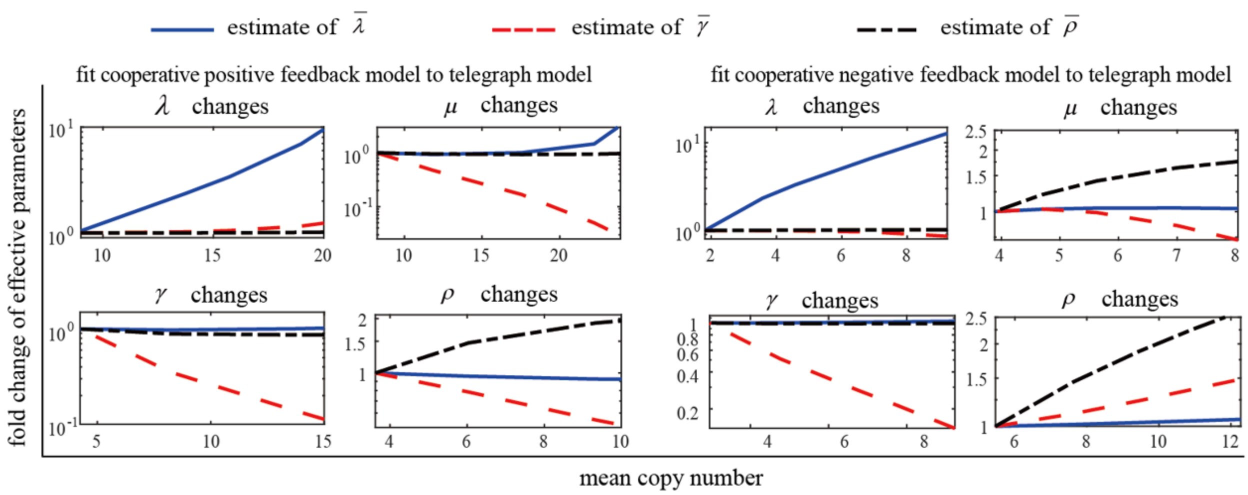


Figure 6: **Variation pattern of effective rate parameters under different induction conditions in cooperative feedback models.** For each positive or negative feedback model with cooperative binding of two proteins, synthetic data of gene product numbers are generated under $625$ parameter sets using the SSA. Tuning a single parameter of cooperative feedback models can generate a series of distributions of gene product numbers, along with different mean expression levels. Fitting these distributions by the telegraph model leads to a series of effective rate parameters $\bar{\lambda}$, $\bar{\gamma}$ and $\bar{\rho}$. Plotting $\bar{\lambda}$, $\bar{\gamma}$ and $\bar{\rho}$ as a function of the corresponding mean expression level reveals how the effective rate parameters vary when the single parameter of cooperative feedback models is tuned. There are general variation patterns of effective rate parameters under different tuning parameters of the cooperative feedback models.

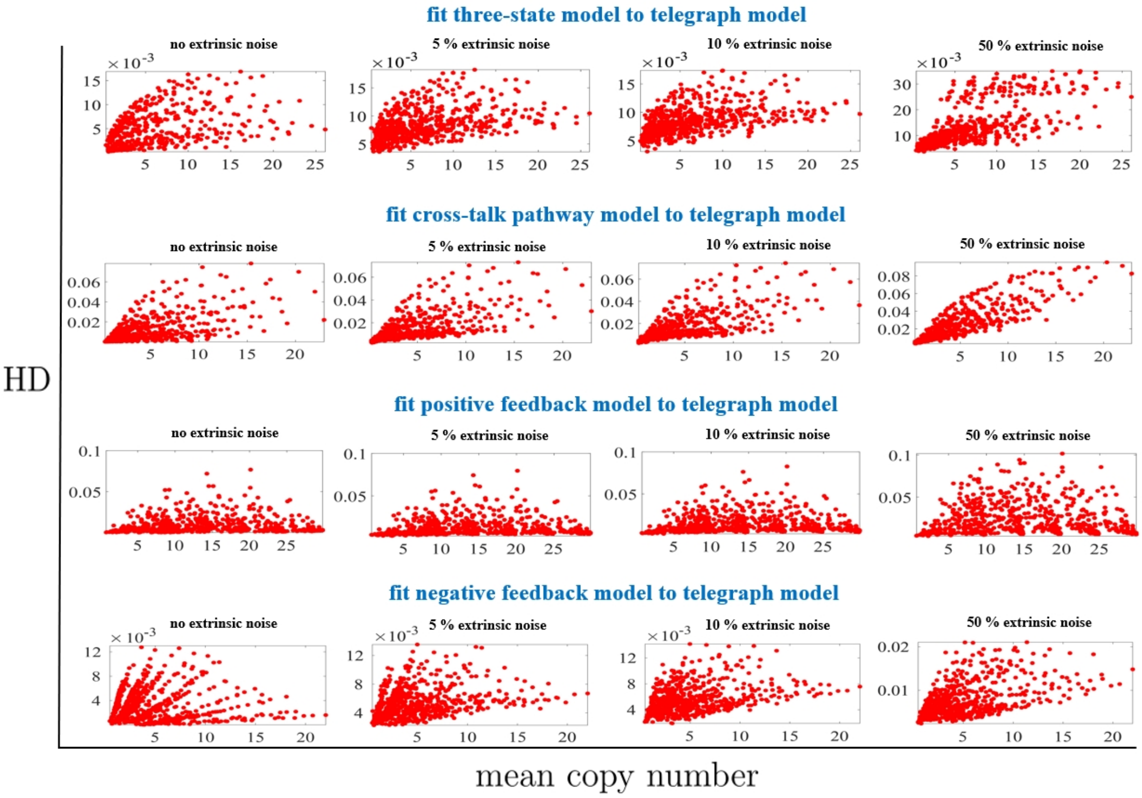


Figure 7: **Fit complex models with extrinsic noise to the telegraph model.** We take into account the extrinsic noise by adding noise to the initiation rate $\rho$ in the three-state, crosstalk pathway, positive feedback and negative feedback models. Here, we reset $\rho$ as a log-normal distributed random variable with its mean being the original value of $\rho$ and standard deviation being equal to 0.05, 0.1, 0.5 of the mean (presenting 5%, 10%, 50% extrinsic noise, respectively). For each of complex models with a extrinsic noise level, synthetic data of gene product numbers are generated under 625 parameter sets using the SSA. In steady state, the Hellinger distance (HD) between the simulated distribution and its telegraph model approximation is shown as a function of the mean expression level for the 625 parameter sets. Under the all tested parameter sets and extrinsic noise levels, the HD is less than 0.035 for the three-state model, less than 0.1 for both cross-talk and positive feedback models, less than 0.021 for the negative feedback model.

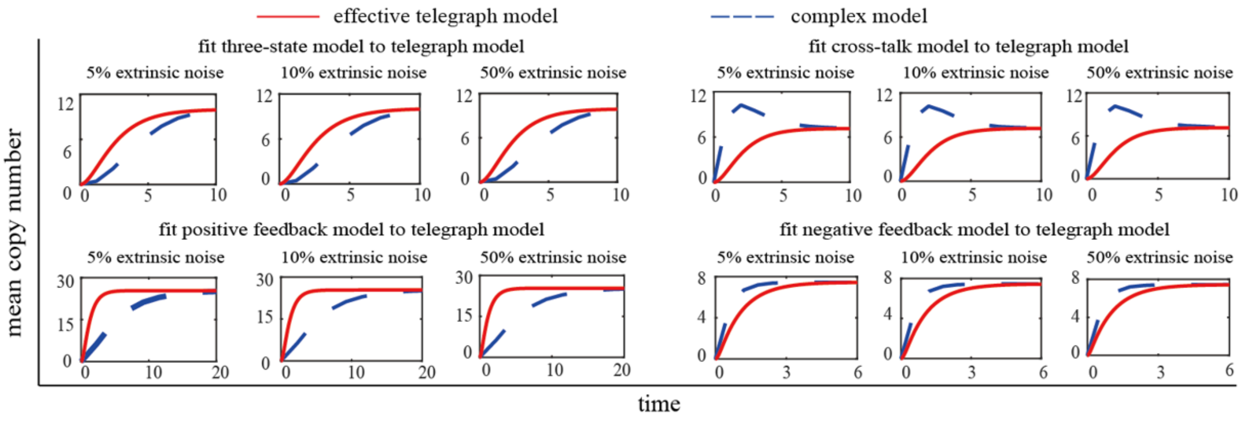


Figure 8: **Distinguishing dynamic differences between complex models under extrinsic noise and their effective telegraph models.** We take into account the extrinsic noise by adding noise to the initiation rate $\rho$ in each of the three-state, cross-talk pathway, positive feedback and negative feedback models. Here, we reset $\rho$ as a log-normal distributed random variable with its mean being the original value of $\rho$ and standard deviation being equal to 0.05, 0.1, 0.5 of the mean (presenting $5\%$, $10\%$, $50\%$ extrinsic noise, respectively). Under the initial condition that the gene is off and there is no gene product molecules in the cell, we generate synthetic data of gene product numbers for each complex model and each extrinsic noise level under $625$ parameter sets using the SSA. We fit the steady-state synthetic data to obtain the effective telegraph models, and plot time-dependent gene products means for both synthetic data and the effective model. For all tested extrinsic noise levels, the three-state and positive feedback models have smaller time-dependent mean curve compared to the effective telegraph model, while the cross-talk pathway and negative feedback models have larger time-dependent mean curve.

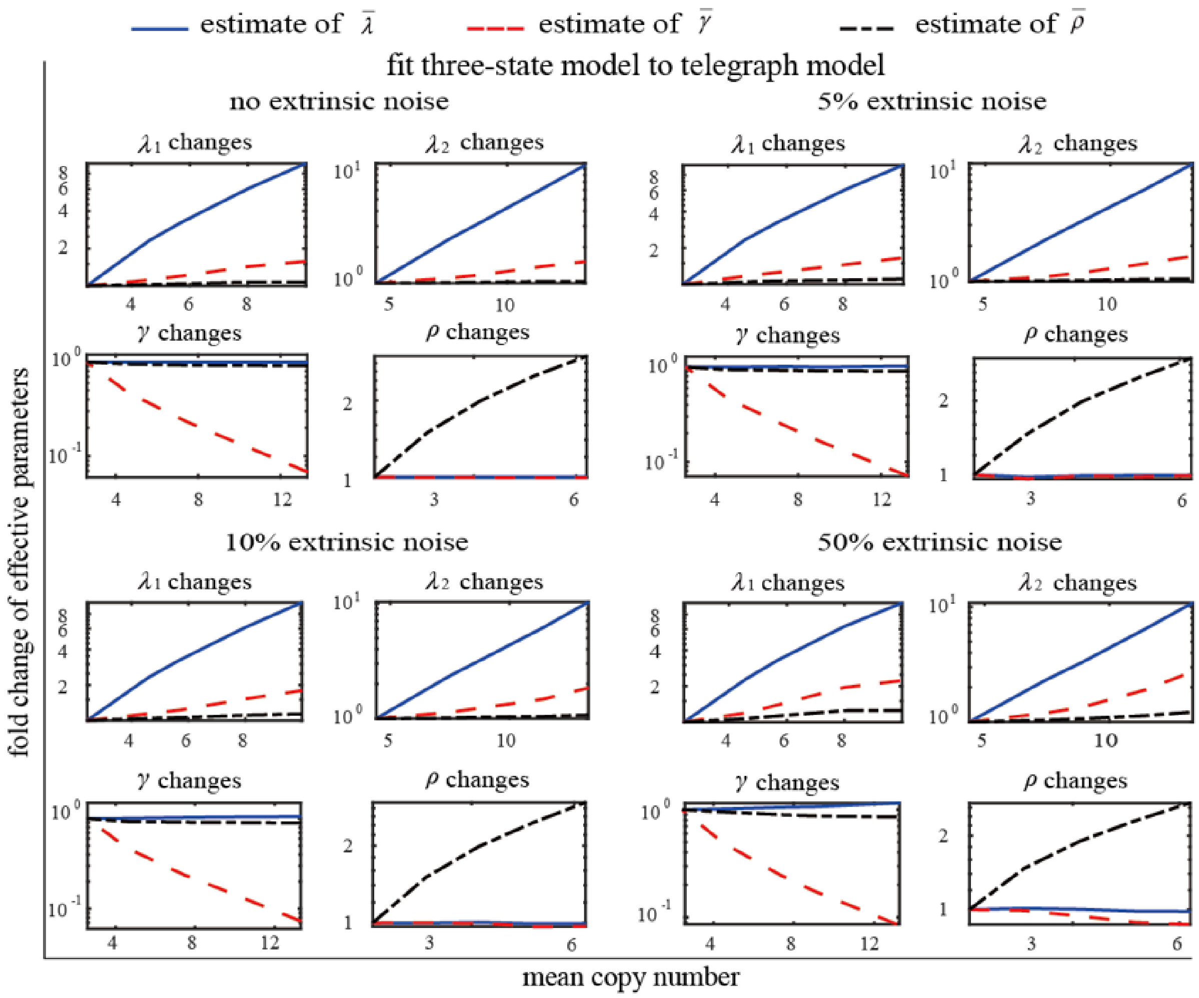


Figure 9: **Variation pattern of effective rate parameters under different induction conditions in the three-state model under extrinsic noise.** We take into account the extrinsic noise by adding noise to the initiation rate $\rho$ in each of the three-state model. Here, we reset $\rho$ as a log-normal distributed random variable with its mean being the original value of $\rho$ and standard deviation being equal to 0.05, 0.1, 0.5 of the mean (presenting 5%, 10%, 50% extrinsic noise, respectively). Tuning a single parameter of the three-state model under each extrinsic noise level can generate a series of distributions of gene product numbers, along with different mean expression levels. Fitting these distributions by the telegraph model leads to a series of effective rate parameters $\bar{\lambda}, \bar{\gamma}$ and $\bar{\rho}$. Plotting $\bar{\lambda}, \bar{\gamma}$ and $\bar{\rho}$ as a function of the corresponding mean expression level reveals how the effective rate parameters vary when the single parameter of the three-state model is tuned. There are general variation patterns of effective rates except for turning $\lambda_1$ (or $\lambda_2$) under 50% extrinsic noise.

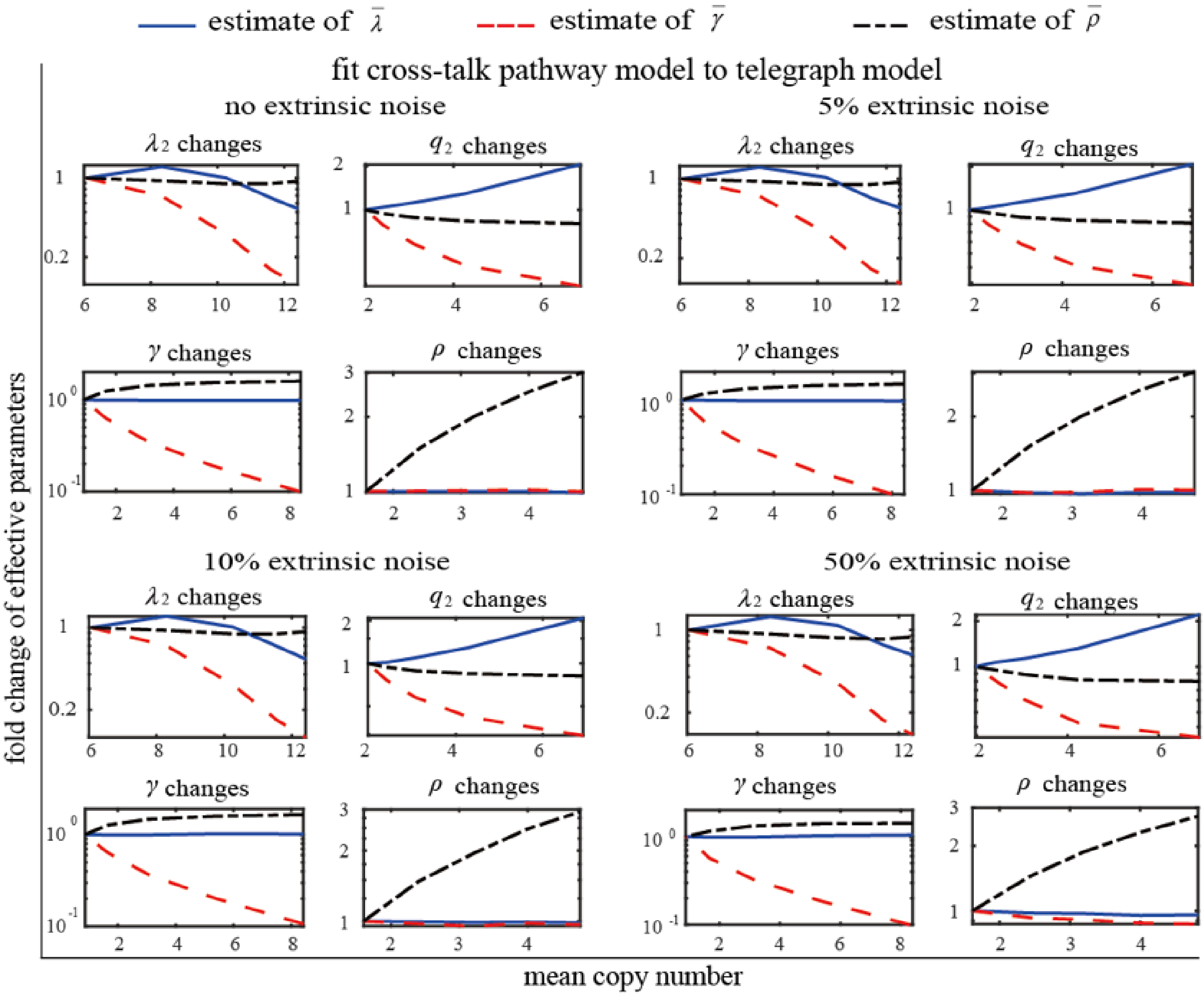


Figure 10: **Variation pattern of effective rate parameters under different induction conditions in the cross-talk pathway model under extrinsic noise.** We take into account the extrinsic noise by adding noise to the initiation rate $\rho$ in each of the cross-talk pathway. Here, we reset $\rho$ as a log-normal distributed random variable with its mean being the original value of $\rho$ and standard deviation being equal to 0.05, 0.1, 0.5 of the mean (presenting 5%, 10%, 50% extrinsic noise, respectively). Tuning a single parameter of the cross-talk pathway model under each extrinsic noise level can generate a series of distributions of gene product numbers, along with different mean expression levels. Fitting these distributions by the telegraph model leads to a series of effective rate parameters $\bar{\lambda}$, $\bar{\gamma}$ and $\bar{\rho}$. Plotting $\bar{\lambda}$, $\bar{\gamma}$ and $\bar{\rho}$ as a function of the corresponding mean expression level reveals how the effective rate parameters vary when the single parameter of cross-talk pathway model is tuned. There are general variation patterns of effective rates under different extrinsic noise levels.

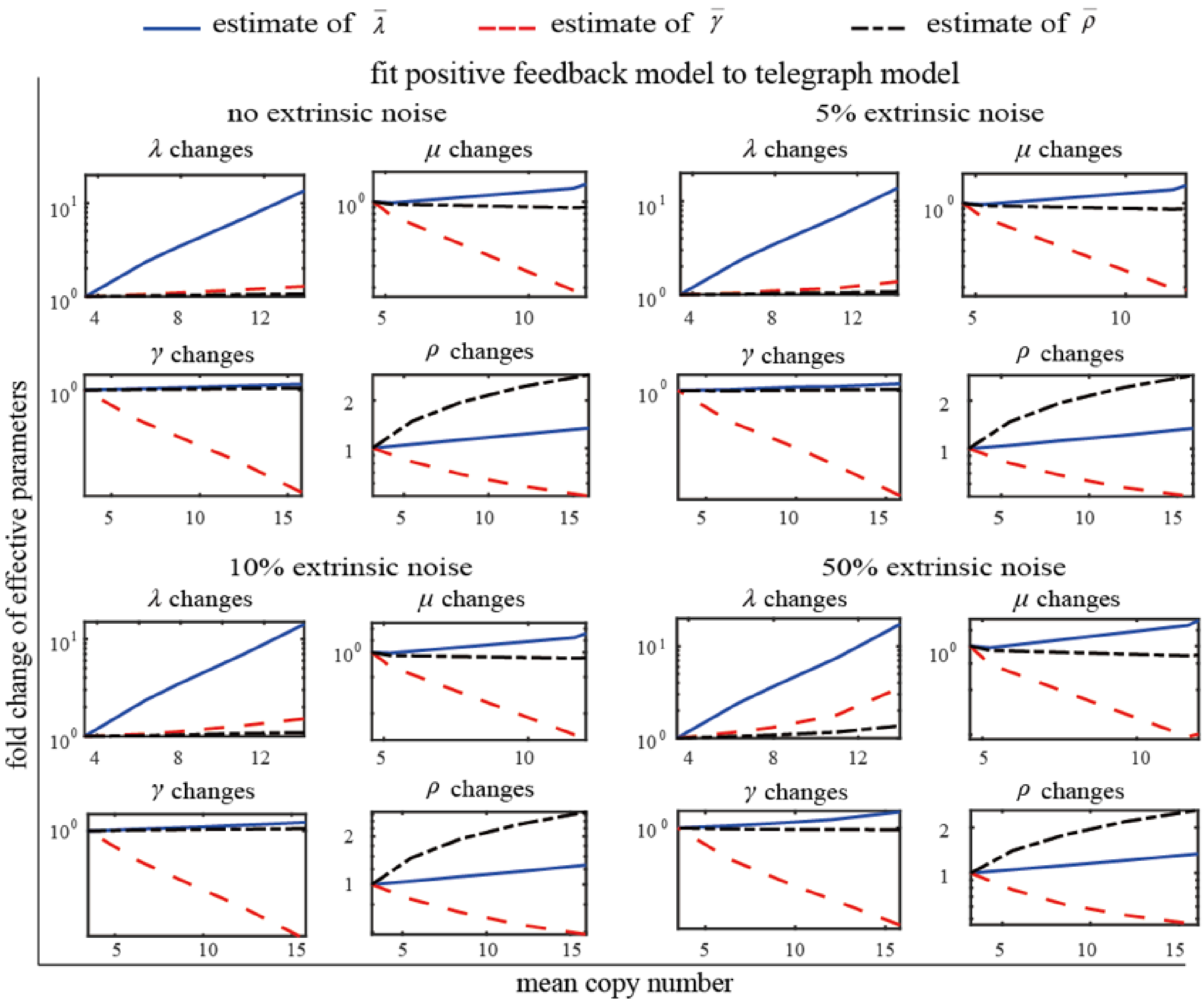


Figure 11: **Variation pattern of effective rate parameters under different induction conditions in the positive feedback model under extrinsic noise.** We take into account the extrinsic noise by adding noise to the initiation rate $\rho$ in each of the positive feedback model. Here, we reset $\rho$ as a log-normal distributed random variable with its mean being the original value of $\rho$ and standard deviation being equal to 0.05, 0.1, 0.5 of the mean (presenting 5%, 10%, 50% extrinsic noise, respectively). Tuning a single parameter of the positive feedback model under each extrinsic noise level can generate a series of distributions of gene product numbers, along with different mean expression levels. Fitting these distributions by the telegraph model leads to a series of effective rate parameters $\bar{\lambda}$, $\bar{\gamma}$ and $\bar{\rho}$. Plotting $\bar{\lambda}$, $\bar{\gamma}$ and $\bar{\rho}$ as a function of the corresponding mean expression level reveals how the effective rate parameters vary when the single parameter of the positive feedback model is tuned. There are general variation patterns of effective rates except for turning $\lambda$ under 50% extrinsic noise.

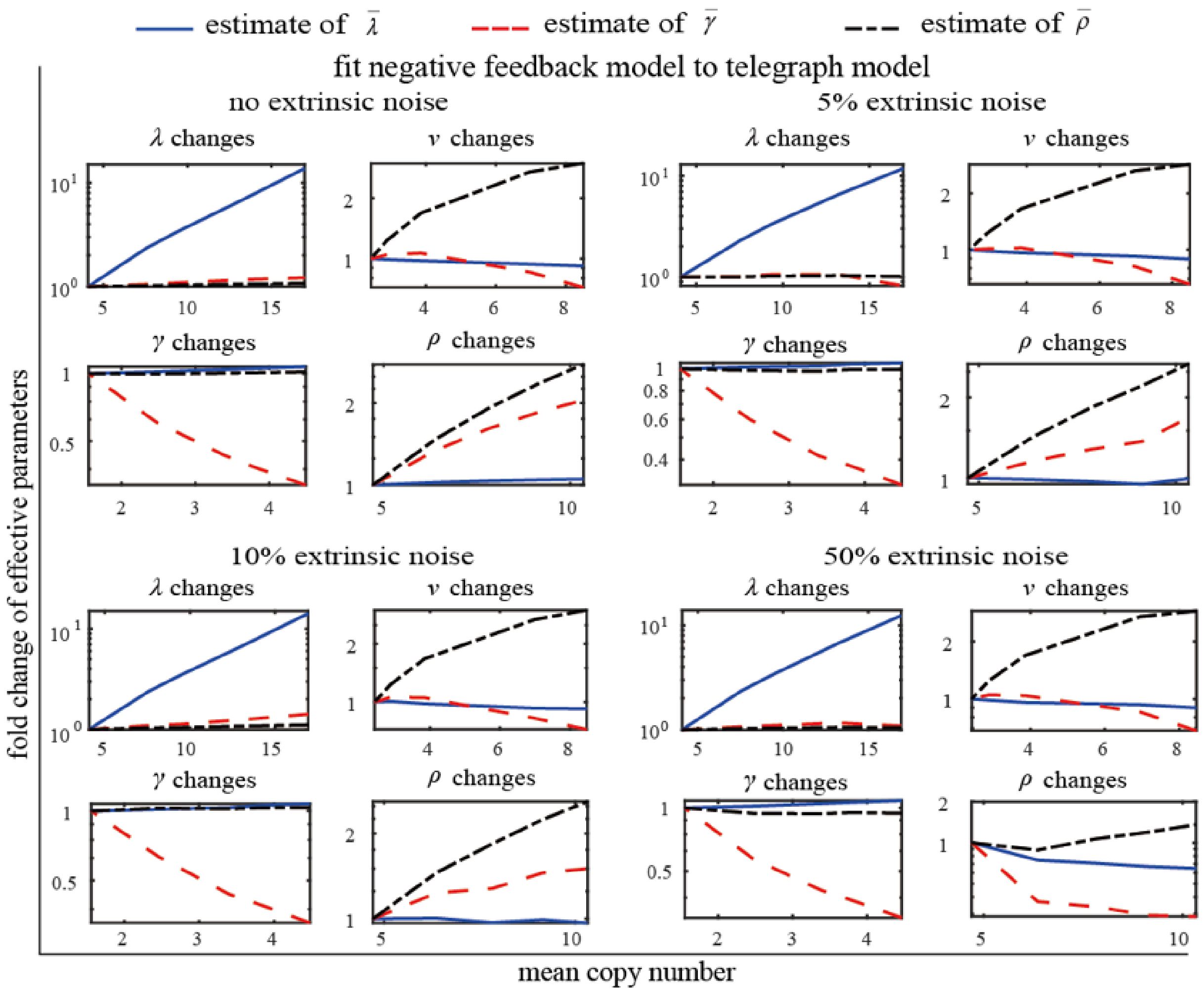


Figure 12: **Variation pattern of effective rate parameters under different induction conditions in the negative feedback model under extrinsic noise.** We take into account the extrinsic noise by adding noise to the initiation rate $\rho$ in each of the negative feedback model. Here, we reset $\rho$ as a log-normal distributed random variable with its mean being the original value of $\rho$ and standard deviation being equal to 0.05, 0.1, 0.5 of the mean (presenting 5%, 10%, 50% extrinsic noise, respectively). Tuning a single parameter of the negative feedback model under each extrinsic noise level can generate a series of distributions of gene product numbers, along with different mean expression levels. Fitting these distributions by the telegraph model leads to a series of effective rate parameters $\bar{\lambda}, \bar{\gamma}$ and $\bar{\rho}$. Plotting $\bar{\lambda}, \bar{\gamma}$ and $\bar{\rho}$ as a function of the corresponding mean expression level reveals how the effective rate parameters vary when the single parameter of the negative feedback model is tuned. There are general variation patterns of effective rates except for turning $\rho$ under 50% extrinsic noise.
